# Clarifying the priming requirement and activator specificity for the NLRP3 inflammasome in human keratinocytes in vitro

**DOI:** 10.1101/2025.08.13.669639

**Authors:** Pritisha Rozario, Ying Shiang Lim, Shirley Suet Lee Ding, Muhammad Jasrie Firdaus, Stephen Wearne, Wong Han Siang Brandon, Rae Chua, Kim S Robinson, Julie Tung Sem Chu, Liu Meng, Shan Sophie Carrie Cai, Sern Ting Eugene Tan, Soon Keong Wee, Mart Matthias Lamers, Navin Kumar Verma, Xia Yun, Eric Peng Huat Yap, John E. A Common, Franklin Zhong

**Affiliations:** Lee Kong Chian School of Medicine, Nanyang Technological University Singapore, 308232, Singapore; A*STAR Skin Research Laboratories (A*SRL), Agency for Science, Technology and Research (A*STAR) Skin Research Labs, 138648, Singapore; Interdisciplinary Graduate Programme, NTU Institute for Health Technologies (HealthTech NTU), Nanyang Technological University, Singapore; The Skin Innate Immunity and Inflammatory Disease Lab, Skin Research Centre, Department of Hull York Medical School, University of York, York Y010 5DD, UK; Programme in Emerging Infectious Diseases, Duke NUS Medical School, Singapore, Singapore; Skin Research Institute of Singapore, 308232, Singapore; Translational and Clinical Research Institute, Newcastle University, Newcastle University, Framlington Place, Newcastle upon Tyne NE2 4HH, UK; Department of Biomedical Sciences, University of Antwerp, Universiteitsplein 12610 Antwerpen, Belgium; National Skin Centre, National Health Group, 308205, Singapore

## Abstract

While NLRP3 has been extensively studied in myeloid cells, its existence and regulation in epithelial cells, including keratinocytes, are unclear. In fact, whether human keratinocytes express a functional NLRP3 inflammasome at all remains a matter of debate in the inflammasome field. Here, we provide additional evidence that NLRP3 is repressed in human keratinocytes cultured under non-inflammatory conditions but can be sharply induced by interferon-γ (IFNγ)—but not lipopolysaccharide (LPS). In this IFNγ-primed state, not all established NLRP3 activators are specific to NLRP3. We report that nigericin-driven keratinocyte pyroptosis occurs via both NLRP1 and NLRP3, whereas *Staphylococcus aureus* α-hemolysin (Hla) exclusively and nonredundantly activates NLRP3, even though both require K+ efflux. Furthermore, in the presence of T cells, certain virulent *S. aureus* strains can cause NLRP3-dependent pyroptotic death in keratinocytes *in vitro* through the cooperative actions of superantigens (SAgs) and Hla. In summary, our findings establish the strict inducibility and functional relevance of the NLRP3 inflammasome in non-myeloid, epithelial cells in vitro. These results resolve conflicting reports and position keratinocytes as a context-specific, non-hematopoietic cellular model for studying NLRP3 activation in host-microbe interactions at barrier tissues.

**KEY FINDINGS:**

1. Additional evidence that NLRP3 is absent in resting, nonstimulated human keratinocytes in vitro
2. IFNγ, but not LPS, is a potent ‘priming’ signal for NLRP3 in human keratinocytes in vitro
3. In IFNγ-primed keratinocytes, *S. aureus* α-hemolysin (Hla) selectively activates NLRP3, whereas nigericin activates both NLRP1 and NLRP3 in vitro
4. SAg and Hla kill keratinocytes via NLRP3-driven pyroptosis in the presence of T cells in vitro

**GRAPHICAL ABSTRACT:** 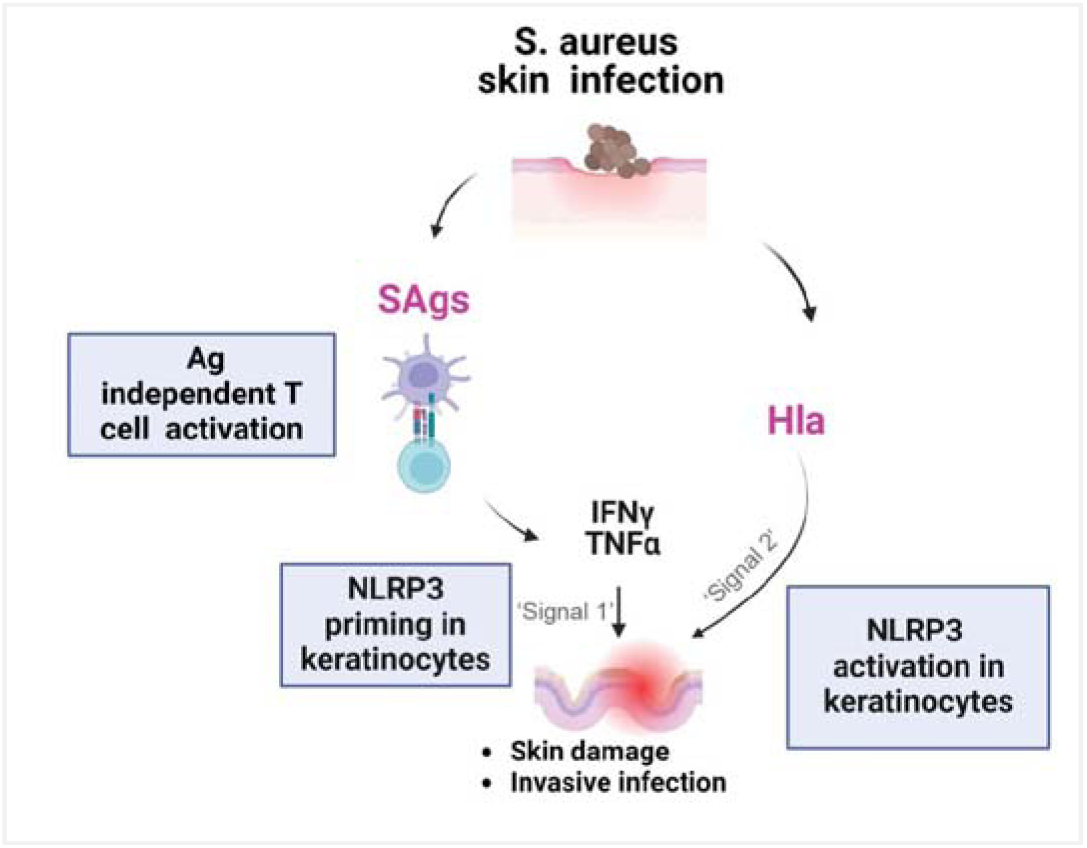

## INTRODUCTION

The inflammasome is a multiprotein complex that plays a critical role in the innate immune system of vertebrates by detecting pathogenic microorganisms and stress signals. In humans, the inflammasome is crucial for maintaining immune homeostasis and protecting against infections, injuries, malignancies and metabolic stress (1–3). At the molecular level, the inflammasome complex is assembled by a sensor protein in the NLR family. Upon detecting microbial infections or cellular damage, NLR proteins oligomerize and recruit adaptor protein ASC and activate caspase-1. Activated caspase-1 processes pro-inflammatory cytokines such as IL-1β and IL-18, and pore-forming protein gasdermin D (GSDMD) into their active forms. Mature IL-1 cytokines are then secreted through GSDMD pores. This sequence of events result in an immunogenic and lytic form of cell death termed pyroptosis (4–8). Pyroptosis can help eliminate infected or damaged cells, prevent the spread of intracellular pathogens and prime the adaptive immune response (9). On the other hand, certain pathogen-encoded virulence factors, especially those from extracellular pathogens can hijack the inflammasome complex to damage host tissues to aid infection or dissemination (10–12). Aberrant and excessive inflammasome activation also contributes to the pathogenesis of many auto-inflammatory and auto-immune disorders (9, 13, 14).

NLRP3 is the most extensively studied inflammasome sensor protein. It is highly expressed in cells of the hematopoietic lineage in mice and humans, particularly monocytes, dendritic cells and macrophages and microglial cells (15). Thus, most in vitro studies on NLRP3 employ murine bone marrow derived macrophages (BMDMs) or human leukemic cell lines, such as THP-1 or U-937. In vitro experiments using these cell types support a two-step model of NLRP3 function. First, a ‘priming’ step (Signal 1), typically involving a Toll-Like Receptor ligand, such as LPS and PAM3CSK, is needed to stimulate NLRP3 mRNA expression. Certain inflammatory cytokines, including TNF, IFNγ, IL-36, IL-17 have also been shown to effectively ‘prime’ NLRP3 expression(16–19). NLRP3 inflammasome can then be triggered by a wide variety of activating molecules such as bacterial toxins (Signal 2) via either a K+ efflux dependent or independent manner (20, 21).

In contrast, the regulation of NLRP3 in non-immune cell types is far less understood. For instance, the field is divided over whether NLRP3 is expressed in human skin keratinocytes, the major cell type of the outer epidermal layer of the skin and likely the first cell type to encounter skin resident and invasive microbes. On one hand, numerous groups have reported constitutive expression of the NLRP3 protein cultured keratinocytes (or related cell lines, such as HaCaT) in vitro and human skin biopsies based on immunoblotting and immunostaining (22–25). Certain pathogen-associated molecular patterns (PAMPs) and toxins, including double-stranded RNA, have been shown to directly activate NLRP3 within keratinocytes, implicating keratinocyte-intrinsic NLRP3 in diverse skin diseases(26, 27) .

In direct conflict with these results, recent studies showed that human keratinocytes cultured under ‘resting’, non-inflammatory conditions predominantly use an alternative inflammasome sensor, NLRP1 to initiate pyroptosis (28–30) (31–33). In fact, a number of groups including ours reported that *NLRP3* mRNA is undetectable in healthy human skin and keratinocytes (31–33). However, these data could not rule out that NLRP3 is expressed in rare subtypes of keratinocytes, or can be induced by specific inflammatory cues. The uncertainty arising from these conflicting results in the literature obstructs clear assessment of the therapeutic potential of NLRP3 inhibitors for skin diseases.

In this study, we aim to clarify the existence and regulation of the NLRP3 inflammasome in human keratinocytes in vitro. We provide additional proof that *NLRP3* mRNA is absent in human keratinocytes cultured under non-inflammatory conditions and nonlesional human skin biopsies. Using *known* NLRP3 priming and activating molecules, we then define the optimal conditions to elicit NLRP3 functionality in human keratinocytes: instead of TLR ligands such as LPS and PAM3CSK, IFNγ is found to be the most effective priming signal to upregulate NLRP3 mRNA in human keratinocytes. In addition, *established* NLRP3 activators in fact differ in their abilities in distinguishing NLRP3 vs NLRP1, with the canonical NLRP3 activator nigericin inducing NLRP1 and NLRP3 simultaneously and alpha-hemolysin (Hla) from *Staphylococcus aureus* (*S. aureus*) exclusively activating NLRP3. Furthermore, we provide preliminary in vitro evidence for a pathway where virulent *S. aureus* strains deploy a combination of two known exotoxins-superantigens (SAg) and Hla to kill human keratinocytes via T-cell initiated paracrine IFNγ priming and NLRP3-driven pyroptosis. By demonstrating the unexpectedly stringent inducibility of NLRP3 mRNA and delineating the signals required to specifically activate human keratinocyte-intrinsic NLRP3 inflammasome, our study position human keratinocytes as an alternative model to investigate endogenous NLRP3 biology in non-immune cell types.

## RESULTS

### NLRP3 expression is repressed in resting keratinocytes and sharply induced by IFN**γ**

Our study is designed to address three questions (Figure 1A): 1) Do human keratinocytes express NLRP3? 2) If so, is NLRP3 regulated differently in keratinocytes from monocytes and macrophages? 3) Can known NLRP3 activators discriminate between NLRP3 and NLRP1 when both are present? To address the first question, we queried existing literature that demonstrated NLRP3 expression in human skin (22–25, 34). Nearly all the studies relied on antibody-based methods, such as immunohistochemistry (IHC), immunofluorescence (IF) and Western blotting (WB) to detect NLRP3. Several studies also used qPCR to measure changes of the NLRP3 mRNA levels. We reasoned that in the absence of genetic knockout controls, it is not possible to evaluate the expression of NLRP3 based on antibody-based methods alone. Similarly, the absolute abundance of the NLRP3 mRNA cannot be accurately measured without in vitro transcribed *NLRP3* mRNA as a spike-in control. These controls are rarely included in published studies.

**Figure 1.**
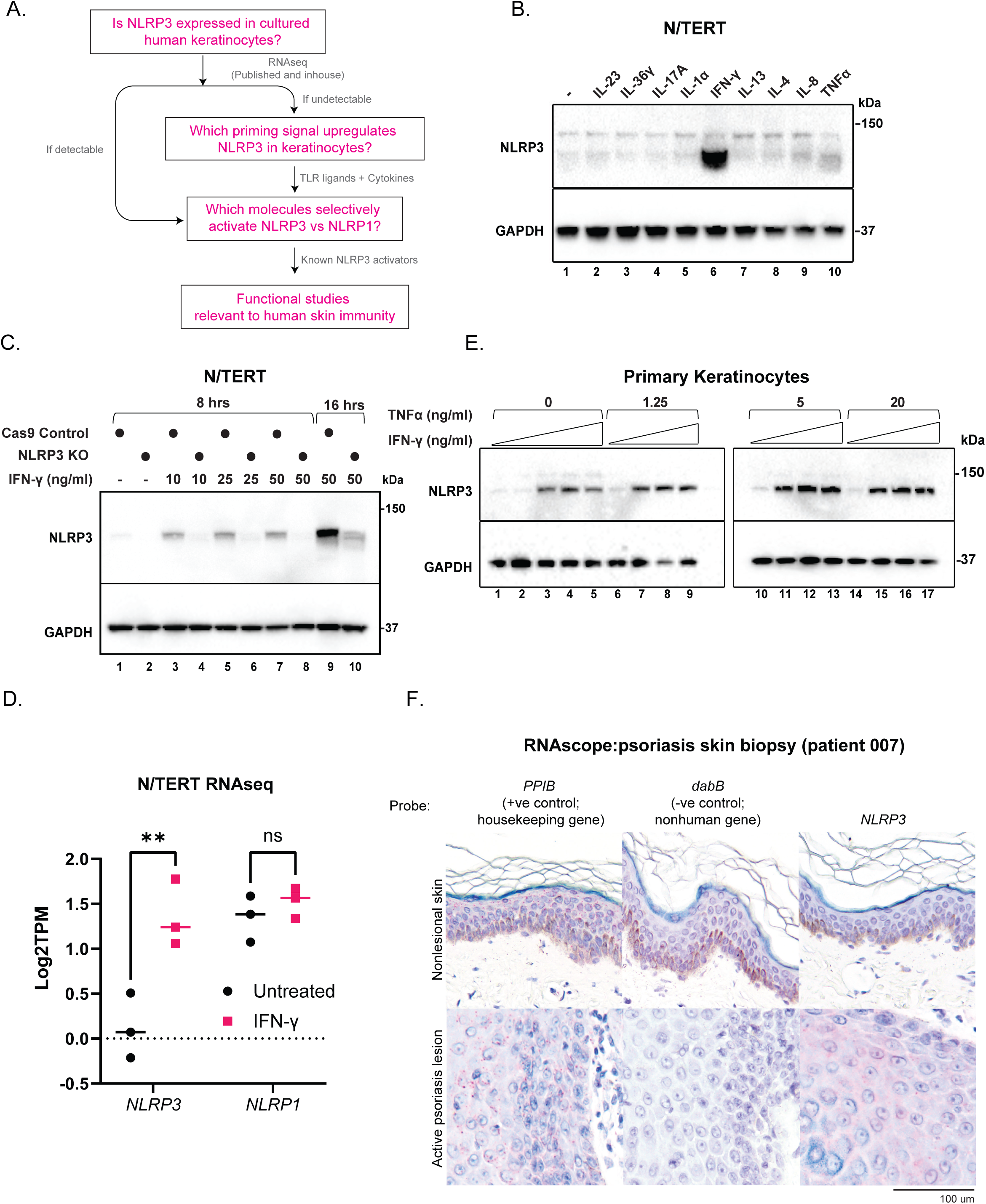
IFNγ is the most robust NLRP3 priming signal in immortalized and primary keratinocytes. A. Summary of study design B. Wild-type (WT) N/TERT upon 16 hrs stimulation with various cytokines at 50 ng/ml. C. Immunoblot of NLRP3 and GAPDH in Cas9 control and NLRP3 KO N/TERT upon stimulation with increasing concentrations of IFNγ for either 8 hrs or 16 hrs. D. Log2 transcripts per kilobase million (TPM) levels of NLRP3 and NLRP1 in untreated vs IFNγ (50 ng/ml) stimulated WT N/TERT cells, treated for 16 hrs. E. Immunoblots of NLRP3 and GAPDH of human primary keratinocytes stimulated with increasing concentrations of IFNγ and TNFα for 16 hrs. F. RNAscope staining of FFPE sections patient skin biopsies. The paired lesional and nonlesional biopsies are obtained from the same patient during a single clinical visit. Immunoblots show lysates from one experiment performed at least three independent times. Error bars represent SEM from three biological replicates, where one replicate refers to independent seeding and treatment of cells. Significance values were calculated based two-way ANOVA followed by Sidak’s test for multiple pairwise comparisons (D) ns, non-significant, ** P<0.01

To more quantitatively assess the abundance of NLRP3 in keratinocytes, we turned to published RNAseq datasets released by three major consortia: ENCODE, HPA and GTex, which collectively profiled more than a thousand human skin samples or donor-derived primary keratinocytes(35–37). In all three datasets, gene level quantification was measured by ‘normalized transcripts per million’ (nTPM), which allows for absolute quantification across different samples and sequencing platforms. We also examined HaCaT cells, an immortalized cell line of keratinocyte origin which is commonly used in skin-related studies and has been shown to express NLRP3 in multiple reports. Arguing against the existence of NLRP3 in keratinocytes and healthy skin, *NLRP3* mRNA level is extremely low (nTPM=0.05-0.4) in all skin-related entries in ENCODE, HPA or GTex, including both bulk RNAseq (averaged across >1000 samples in total) and scRNA seq (basal vs differentiated keratinocytes). In fact, *NLRP3* nTPM is even lower than that of the control liver-specific transcript, albumin (*ALB*). By contrast, NLRP1 is detectable in all skin/keratinocytes samples (minimum nTPM>4.5) (Table 1).

**Table 1:** Additional evidence that NLRP3 mRNA is not expressed in unperturbed human skin keratinocytes or HaCaT cells

|  |  |
| --- | --- |
| Data source | ENCODE ENCSR000CPK |
| Cell/tissue origin and assay | **Human primary keratinocytes** (female). polyA plus RNA-seq |
| **Gene quantification/normalized TPM** |  |
| *NLRP3* | **0.05** |
| *NLRP1* | 9.61 |
| *TLR4* | 0.04 |
| *TLR2* | 0.25 |
| *ALB* (albumin), -ve control | 0.01 |
| Data source | HPA RNA-seq tissue data |
| Cell/tissue origin and assay | **Healthy human skin**. Total RNAseq |
| **Gene quantification/normalized TPM** |  |
| *NLRP3* | **0.2** |
| *NLRP1* | 38.3 |
| *TLR4* | 0.9 |
| *TLR2* | 2.6 |
| *ALB* (albumin), -ve control | 0.3 |
| Data source | GTEX RNA-seq tissue data |
| Cell/tissue origin and assay | **Human Skin - Not Sun Exposed (Suprapubic)+Sun Exposed (Lower leg)**.Total RNAseq<br>Skin - Sun Exposed (Lower leg) |
| **Gene quantification/normalized TPM** |  |
| *NLRP3* | **0.3** |
| *NLRP1* | 20 |
| *TLR4* | 3.4 |
| *TLR2* | 3.4 |
| *ALB* (albumin), -ve control | 3.2 |
| Data source | GSE130973. Normalization in Human Protein Atlas |
| Cell/tissue origin and assay | Human skin: **basal keratinocytes**. Single cell sequencing |
| **Gene quantification/normalized TPM** |  |
| *NLRP3* | **0.4** |
| *NLRP1* | 13.6 |
| *TLR4* | 0.4 |
| *TLR2* | 0.8 |
| *ALB* (albumin), -ve control | 0 |
| Data source | GSE130973. Normalization in Human Protein Atlas |
| Cell/tissue origin and assay | Human skin: **suprabasal keratinocytes**. Single cell sequencing |
| **Gene quantification/normalized TPM** |  |
| *NLRP3* | **0.1** |
| *NLRP1* | 20.2 |
| *TLR4* | 0.7 |
| *TLR2* | 1.3 |
| *ALB* (albumin), -ve control | 0.1 |
| Data source | HPA+Cancer Cell Line Encyclopedia |
| Cell/tissue origin and assay | HaCaT. Total RNAseq |
| **Gene quantification/normalized TPM** |  |
| *NLRP3* | 0 |
| *NLRP1* | 4.5 |
| *TLR4* | 0.1 |
| *TLR2* | 6 |
| *ALB* (albumin), -ve control | 1.5 |

Next we asked if NLRP3 can be induced by known TLR ligands or cytokines in keratinocytes. Similar to primary keratinocytes, NLRP3 was hardly detectable in N/TERT cells without any exogenous stimulation (nTPM<1, Figure 1B-D, Figure S1A-B). No upregulation of NLRP3 protein was observed in N/TERT cells with LPS and Pam3CSK4, the most commonly used priming molecules in hematopoietic cells, or type I/III interferons (Figure S1A-B). However, a specific NLRP3 band became detectable by immunoblot after priming with IFNγ, extracellular Poly(I:C) and to a smaller extent, TNFα (Figure 1B, S1A-B), with IFNγ being the strongest (Figure 1B, S1B). Importantly, the specificity of the NLRP3 Western blotting signal was confirmed by NLRP3 knockout (Figure 1C) and the on-target specificity of recombinant IFNγ was confirmed by chemical inhibition of JAK kinases (Figure S1D). Importantly, even though NLRP3 could be detected reliably by immunoblotting, we were unable to find any commercial antibodies that can distinguish wild-type and NLRP3 KO cells by immunofluorescence. Using bulk RNAseq, we confirmed that the induction of NLRP3 by IFNγ occurs at the level of mRNA transcription (Figure 1D), consistent with a ‘priming’ effect. IFNγ also upregulated other inflammasome components, including pro-caspase-1 and pro-IL-18 (Figure S1C), in agreement with published results, but it had no effect on NLRP1 (Figure S1E). It is well known that IFNγ can prime NLRP3 in microglial cells, endothelial cells, macrophages and THP-1 cells, but it is generally less potent than TLR ligands such as LPS and PamCSK4 (38–40). Indeed, equivalent concentrations of IFNγ failed to efficiently prime NLRP3 in human PBMCs in our hands (Figure S2B). Taken together, these results show that human keratinocytes uniquely rely on IFNγ to prime NLRP3 expression. The inability of LPS and Pam3CSK4 to effectively prime NLRP3 in keratinocytes is likely due to the low levels of the corresponding TLRs (Table 1).

Recent work has shown that IFNγ and TNFα, when present together in the extracellular milieu, act synergistically and result in a greater inflammatory response than either cytokine alone (41, 42). Such synergism also holds true for NLRP3 priming, as IFNγ and TNFα together led to greater NLRP3 induction in N/TERT cells (Figure S1E) as well as human primary keratinocytes (Figure 1E) than IFNγ alone.

Next we tested if *NLRP3* mRNA is induced in skin diseases that involve IFNγ-and TNFα-driven inflammation. In paired nonlesional and lesional biopsies obtained from two patients with plaque psoriasis, RNAscope in situ staining revealed that *NLRP3* mRNA is absent in nonlesional skin, but detectable in active psoriatic lesions (Figure 1F, Figure S2A). Although we are not able to quantify the absolute local concentrations of IFNγ and TNFα in archival FFPE sections, both cytokines are well known to be elevated in psoriasis lesions and directly contribute to disease progression(43–45). Thus our results are consistent with the in vitro finding that NLRP3 is repressed in the absence of inflammatory cues but can be sharply induced by specific inflammatory cytokines, especially IFNγ. This phenomenon seems to be specific to human skin, as IFNγ+TNFα failed to induce NLRP3 expression in human nasal epithelial cells or iPSC-derived kidney organoids (Figure S2C-D). Our data suggest a plausible explanation on the conflicting literature regarding NLRP3 in keratinocytes: it is possible that certain keratinocyte growth conditions elicit tonal inflammatory signaling involving the IFNγ-STAT1 pathway, inadvertently priming NLRP3 expression.

### IFN**γ** primed human keratinocytes as a new model to study NLRP3 function in epithelial cells

IFNγ-primed keratinocytes offer a unique non-immune cellular model that expresses all components of the NLRP3 and NLRP1 inflammasome components at the endogenous level. Thus, it offers an ideal system to perform comparative studies of these two inflammasome sensors, which is not possible with immune cell types. This is especially pertinent, given that many potassium ionophores, including the most widely used NLRP3 activator nigericin, also activates NLRP1 via the ribotoxic stress kinase ZAK (46, 47). However, as NLRP3 is not expressed in unprimed keratinocytes, it has not been possible to rigorously test whether this agonism is truly shared when both NLRP1 and NLRP3 are present. We thus tested this possibility in IFNγ+TNF -primed N/TERT cells. Nigericin was indeed observed to activate both sensors, as deleting NLRP1 or NLRP3 partially reduced, but was not sufficient to eliminate nigericin-driven IL-1β secretion (Figure S3A-B). Abrogation of IL-1β secretion and GSDMD cleavage was only observed when primed NLRP1 KO cells were further treated with NLRP3 inhibitor MCC950, or when both NLRP3 and ZAK were inhibited chemically (Figure S3C). In this context, MCC950 had a greater effect than Compound 6p (ZAK), consistent with our previous observation of NLRP3 is more sensitive than NLRP1 towards K+ efflux (Figure S3C).

Although both NLRP1 and NLRP3 can be activated by K+ efflux, we previously reported that NLRP1 is much less tolerant of other ionic flux than NLRP3. For instance, less selective perforation of the plasma membrane by gramicidin and palytoxin activates NLRP3 but fails to activate NLRP1, despite efficient K+ efflux (46). We therefore reasoned that large membrane pores formed by bacterial pore forming toxins would likely selectively activate NLRP3 in primed keratinocytes without concomitantly activating NLRP1. To test this hypothesis, we focused on *S. aureus* alpha-hemolysin (Hla). Hla is a well known NLRP3 activator in murine bone marrow-derived macrophages (BMDMs) and human macrophages and has an established role in the pathogenesis of *S. aureus* infection in various disease models, including the skin (12, 48, 49). Importantly, the structure of Hla pores are >1 nm in diameter, sufficient for the movement of larger molecules than K+ (50). Other bacterial pore forming toxins, including those in the alpha-hemolysin family have also been shown as definitive NLRP3 activators in macrophages(21, 51).

In unprimed N/TERT cells, Hla alone was sufficient to cause cytotoxicity as evidenced by DRAQ7 uptake; however, this was apparent only at a relatively late time point (∼10 hours) (Figure 2A). By contrast, Hla treatment of IFNγ+TNFα stimulated N/TERT led to DRAQ7 uptake (Figure 2B) much more rapidly (∼ 3-4 hours), consistent with pyroptosis. In the presence of MCC950, DRAQ7 uptake was significantly delayed (Figure 2B). MCC950 also abrogated GSDMD cleavage in Hla-treated primed cells, but had no effect on anisomycin (ANS)-or VbP-induced NLRP1 inflammasome activation (Figure S4A). Additionally, genetic deletion of NLRP3, similar to MCC950 treatment, abrogated hallmarks of inflammasome activation induced by Hla in primed N/TERT cells, including ASC speck formation and IL-1β secretion (Figure 2C-E, Figure S4B-C). Hla-driven NLRP3 activation could be partially, but significantly inhibited by extracellular K+ supplementation (Figure S4D), in keeping with its known mechanism of NLRP3 activation in macrophages. Hla does not trigger NLRP1 inflammasome activation in N/TERT cells, as it failed to induce any pyroptosis in the absence of priming or in primed NLRP1 KO cells (Figure 2E, Figure S4B-C). Thus, *S. aureus* Hla functions as a selective NLRP3 activation signal that does not co-activate NLRP1.

**Figure 2.**
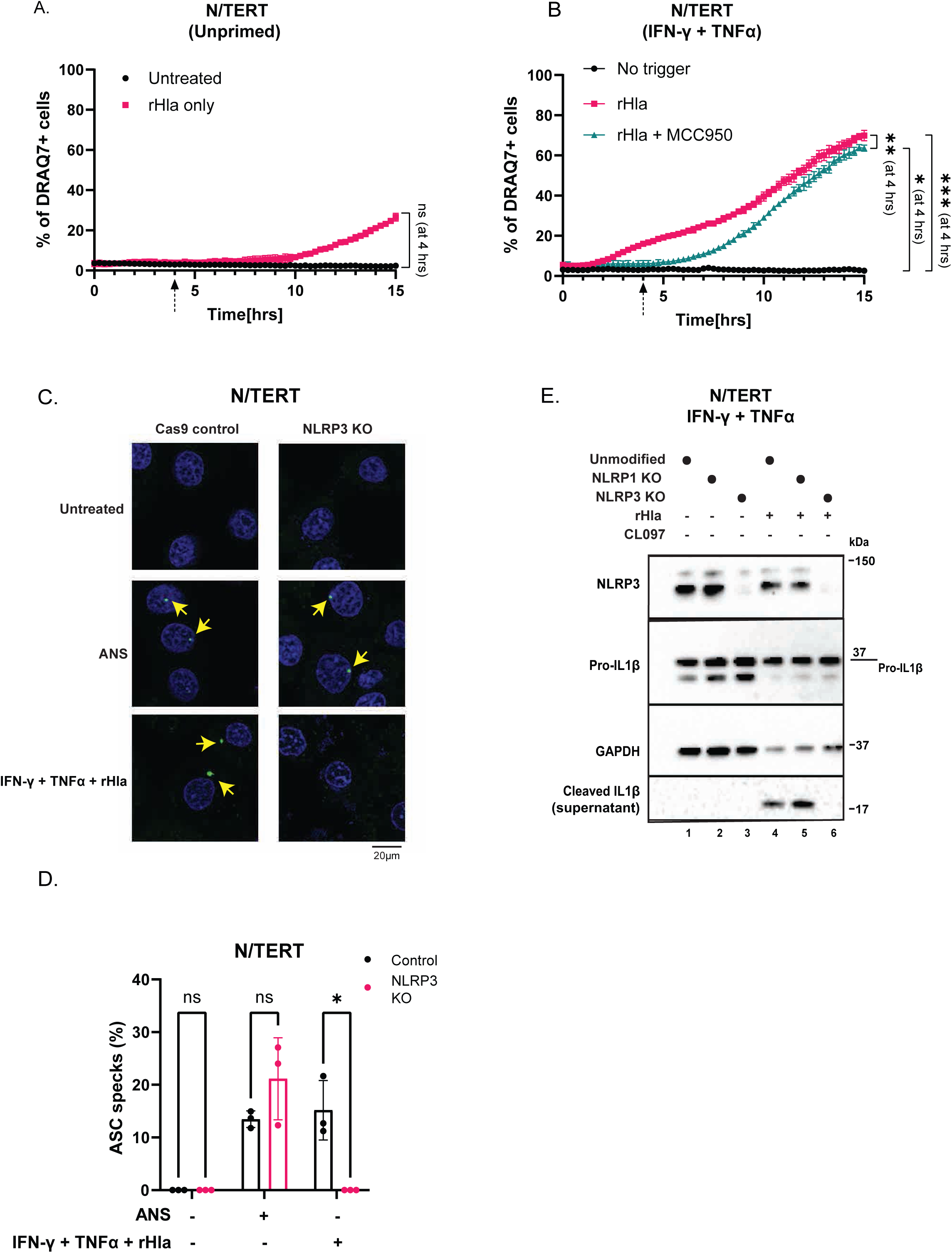
Recombinant Hla activates NLRP3 but not NLRP1 in primed human keratinocytes. A. Quantification of the percentage of DRAQ7 +ve cells in unprimed N/TERTs upon stimulation with recombinant Hla (rHla) (1 μg/mL). B. Quantification of the percentage of DRAQ7 +ve cells N/TERTs ‘primed’ overnight with IFNγ (50 ng/ml) + TNFα (20 ng/ml) followed by rHla (1 μg/mL) +/-MCC950 (5 μM). Cells were imaged at 15 min intervals for 15 hrs upon treatment with rHla. Statistical significance was performed at the 4 hour time points, indicated by arrows. C. Representative confocal microscopy images showing DAPI in blue and anti-ASC immunostaining in green. Images are shown from one experiment and are representative of n = 3 independent experiments; scale bar, 20 μm. Cells were primed overnight and stimulated for 3 hours as indicated in the presence of caspase-1 inhibitor, emricasan (5 μM). Yellow arrows indicate ASC specks. D. Quantification of the ratio ASC specks and the total number of cells scored. E. Immunoblots of NLRP3, pro-IL-1β, cleaved IL-1β and GAPDH (loading control) in WT vs NLRP1 KO vs NLRP3 KO N/TERT following overnight stimulation with IFNγ (50 ng/ml) + TNFα (20 ng/ml) and overnight treatment with rHla (1 μg/mL) Immunoblot shows lysates from one experiment performed three independent times. Error bars represent SEM from three biological replicates, where one replicate refers to independent seeding and treatment of cells. Significance values were calculated based unpaired t test with Welch’s correction at time - 4 hrs (A) ns, non-significant, * P<0.1, ** P<0.01, *** P<0.001

We also sought to validate the functionality of NLRP3 in keratinocytes using an unrelated molecule, CL097, which has been shown to induce NLRP3 assembly in murine BMDMs via ROS, instead of K+ efflux (52, 53). Treatment of IFNγ-primed primary keratinocytes with CL097 indeed resulted in NLRP3-dependent inflammasome activation as measured by IL-1β and GSDMD cleavage (Figure S5A-B). This activation was fully inhibited by the caspase-1 inhibitor Belnacasan and MCC50 (Figure S5A-B). We also confirmed that Hla induces IL-1β in IFNγ+TNF primed primary keratinocytes (Figure S5C), which was abrogated by caspase-1 inhibitor belnacasan but unaffected by necrostatin (Figure S5C). These results confirm that once primed with IFNγ and TNFα, primary keratinocytes and N/TERT cells encode a fully functional NLRP3 inflammasome, which can respond to both K+ efflux dependent and independent activating signals.

Next we addressed the role of NEK7 in NLRP3 and NLRP1 activation. NEK7 was first identified as a crucial interacting partner of NLRP3 in mice, essential for NLRP3 activation in murine bone marrow derived macrophages (54–56). On the contrary, NEK7 appears dispensable for NLRP3 activation in human myeloid cells (57). Recently, NEK7 was also shown to mediate human NLRP1 activation (58). In our hands, deletion of NEK7 did not impair NLRP1 inflammasome activation by anisomycin or VbP (Figure S6A) in unprimed N/TERT cells, arguing against a role of NEK7 in NLRP1 function, at least in N/TERT cells. We observed a reduction of Hla-driven GSDMD and IL-1β cleavage in IFNγ and TNF primed NEK7 KO N/TERT cells, but the magnitude of this reduction was smaller than that of MCC950 or NLRP3 deletion (Figure S6B, Figure 2E). These results are in broad agreement with those reported in Schmacke et al (57): similar to human macrophages, NEK7 is partially involved in, and not strictly essential for NLRP3 inflammasome activation in primed N/TERT cells. Future studies are required to further delineate the precise contribution of NEK7 to human NLRP3 activation.

### Hla is the predominant NLRP3 activating *S. aureus* exotoxin

Hla is among the many *S. aureus* exotoxins that are known to activate NLRP3 in human or murine macrophages (59)(60). NLRP3 has also been shown to play an important role in *S. aureus* infection in mice, where inflammasome components are not expressed in epidermal keratinocytes(61). The role of *keratinocyte-intrinsic* NLRP3 inflammasome in *S. aureus* human skin infection, and the specific contribution of Hla, is less clear. We therefore investigated whether Hla is the predominant or a redundant factor used by *S. aureus* to activate keratinocyte-specific NLRP3. First, we established a small collection of *S. aureus* strains, including two laboratory adapted strains (SH1000, Rosenbach ATCC29213), one MRSA strain (USA300) and four previously uncharacterized strains isolated from distinct body sites of healthy donors (Figure 3B). A common skin commensal species, *S. epidermidis* was included as a control. Prior to stationary phase growth, the amount of Hla secretion by all seven strains was measured using semi-quantitative anti-Hla immunoblot and normalized to bacterial count (CFU/ml) (Figure 3A, S7A-C). In parallel, the sterile filtrates of all seven strains were used to stimulate primed N/TERT ASC-GFP reporter cells at various dilutions (Figure 3A). The ability of each *S. aureus* strain to activate NLRP3, as measured by ASC-GFP speck formation, correlated closely with the amount of Hla it secretes (Figure 3B-C, S7E-G). We found the strains differ widely in terms of Hla expression. Hla^high^ strains USA300, SH1000 and 0857 (defined as >50 μg Hla / million CFU) were on average >3 fold more effective at inducing ASC-GFP specks in primed ASC-GFP reporter N/TERT cells than Hla^low^ strains (<20 μg Hla / million CFU). Thus, the ability for *S.aureus* strains to stimulate NLRP3 in N/TERT ASC-GFP reporter cells correlates with the Hla level of the strain.

**Figure 3.**
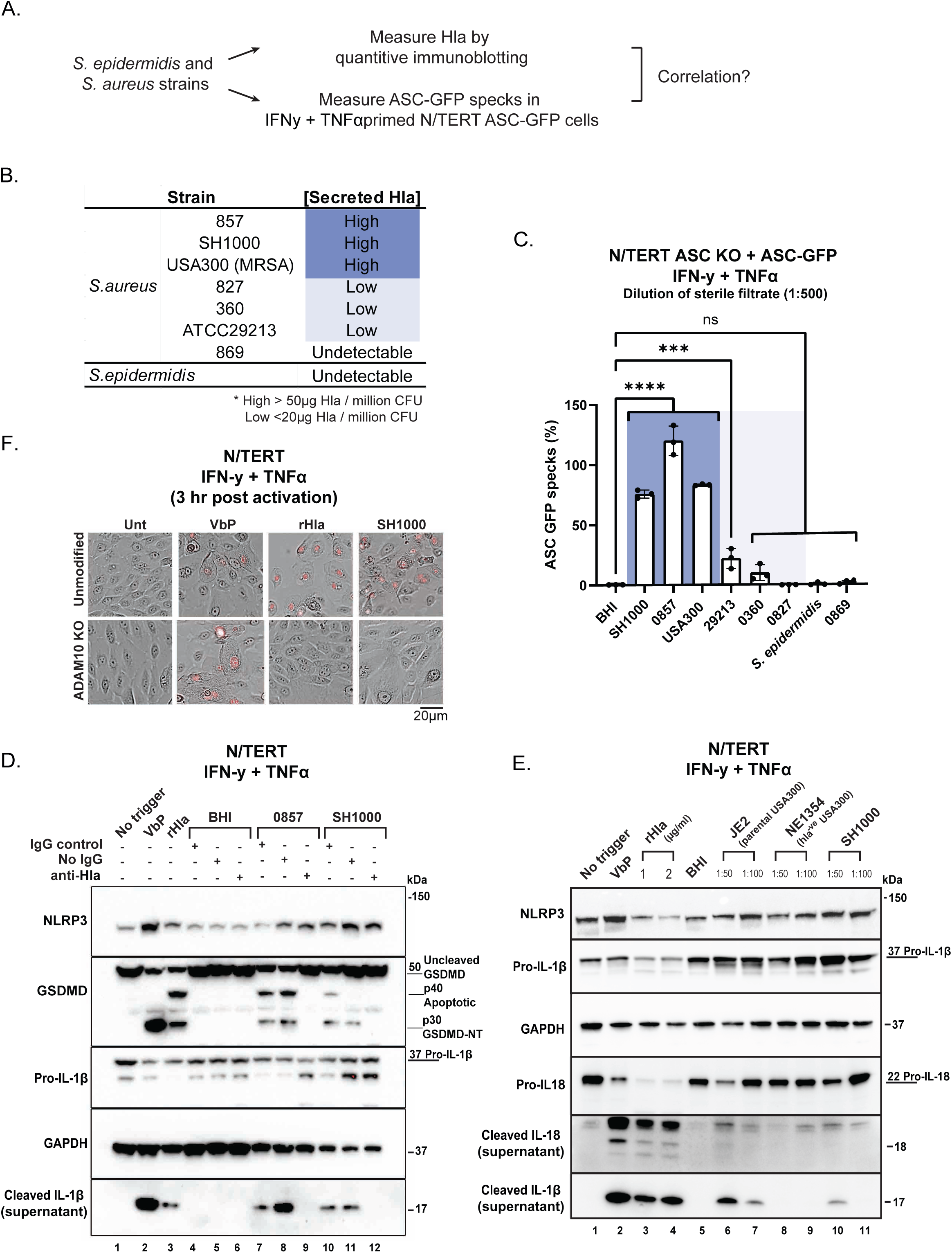
Hla is the predominant, if not the only *S. aureus* exotoxin that activates keratinocyte-intrinsic NLRP3. A. Schematic workflow on measuring Hla secretion from *S. epidermidis* and various *S. aureus* strains and assessing their ability to trigger NLRP3 inflammasome activation in the N/TERT ASC KO + ASC-GFP reporter cells. B. Strains of *S. aureus* tested and Hla levels. Strains were categorized as “High” (Hla > 50 μg/million CFU), “Low” (Hla < 20 μg/million CFU), or “Undetectable” (no quantifiable Hla). C. Percentage of ASC-GFP specks forming cells in N/TERT ASC KO + ASC-GFP cells stimulated overnight with IFNγ (50 ng/ml) + TNFα (20 ng/ml) followed by 3 hrs activation with various sterile filtrates in the presence of emricasan (5 μM). All sterile filtrates were used at 1:500 dilution except USA300 (1:50) due to its low CFU count. Color shades correspond to Hla levels those used in B. D. Immunoblots of NLRP3, GSDMD (full length and cleaved), pro-IL-1β, cleaved IL-1β and GAPDH (loading control) of WT N/TERT stimulated overnight with IFNγ (50 ng/ml) + TNFα (20 ng/ml) followed by treatment with Vbp (3 μM), rHla (1 μg/ml), BHI, 0857 or SH1000 +/-IgG control or anti-α hemolysin. BHI and the respective bacterial sterile filtrates were used at 1:1000 dilution. E. Immunoblots of NLRP3, pro-IL-1β, cleaved IL-1β, pro-IL-18 and cleaved IL-18 and GAPDH (loading control) of WT N/TERT stimulated overnight with IFNγ (50 ng/ml) + TNFα (20 ng/ml) followed by treatment with Vbp (3 μM), rHla (1 μg/ml and 2 μg/ml), BHI media, sterile filtrates of JE2 (Genotype:USA300), NE1354 (Hla KO) or SH1000. BHI was used at 1:50 dilution and the respective bacterial sterile filtrates were used at 1:50 or 1:1001000 dilutions as stated. F. Brightfield microscopy images of WT vs ADAM10 KO N/TERTs stimulated overnight with IFNγ (50 ng/ml) + TNFα (20 ng/ml) followed by activation with Vbp (3 μM), rHla (1 μg/ml) or SH1000 (1:1000) with DRAQ7. Images were obtained 3 hrs post activation. Images are shown from one experiment and are representative of n = 3 independent experiments; scale bar, 20 μm. Immunoblot shows lysates from one experiment performed three independent times. Error bars represent SEM from three biological replicates, where one replicate refers to independent seeding and treatment of cells. Significance values were calculated based on one-way ANOVA followed by Dunnett’s test (C) ns, non-significant; ***P < 0.001; ****P < 0.0001.

To functinally assess the contribution of Hla, we measured the effect of a neutralizing Hla antibody on NLRP3 activation induced by Hla^high^ strains 0857 and SH1000. As compared to an isotype control IgG, pre-incubation with anti-Hla antibody eliminated GSDMD and IL-1β p17 cleavage in primed N/TERT cells (Figure 3D).

To more definitively test the necessity of Hla, we assessed NLRP3 activation induced by Hla knockout MRSA strain (NE1354) and Hla-sufficient parental strain (JE2). Hla^high^ SH1000 was included as a positive control (Figure S7D). Relative to the parental strain JE2 and Hla^high^ strain SH1000, Hla-deficient NE1354 failed to induce IL-1β p17 and IL-18 cleavage in primed N/TERT cells (Figure 3E).

We next used CRISPR/Cas9 to knock out the host Hla receptor ADAM10 (Figure S8B) (62). ADAM10 KO abrogated key features of pyroptosis including DRAQ7 uptake, GSDMD cleavage and IL-1β p17 in primed N/TERT cells treated with Hla or SH1000 filtrate (Figure 3F, S8C). These results prove that of all the many secreted *S. aureu*s exotoxins, Hla is the prominent, if not the only one that activates keratinocyte-specific NLRP3 inflammasome. Other *S. aureus* exotoxins which have previously been shown to activate NLRP3 may have evolved to target cell types other than keratinocytes (59, 63).

### SAg-activated T cells are a likely source of the IFN**γ** and TNF**α** priming signal during *S. aureus* skin infection

IFNγ is a pleiotropic cytokine that plays a crucial role in antifungal and anti-mycobacterial defense, but its function in anti-staphylococcal immunity is less defined. We thus investigated the potential source of endogenous IFNγ and/or TNFα in *S. aureus* infection. *S. aureus* secrete a class of structurally related exotoxins collectively termed ‘super-antigens’ (SAgs) (64–66). Biochemically, SAgs crosslink major histocompatibility complex (MHC) class II molecules on antigen-presenting cells with T-cell receptors, bypassing the need for cognate antigens. This allows SAgs to activate a large fraction of T cells, resulting in massive T cell proliferation and release of proinflammatory cytokines including IFNγ, TNFα and IL2, leading to toxic shock *in vivo* (67). SAgs are highly polymorphic among *S. aureus* strains, with more than 20 characterized to date. Some strains do not encode any SAgs, with others encoding several. The presence of SAgs correlates with virulence of the strains in certain contexts (68, 69).

We hypothesized that SAg-driven T cell activation provides the endogenous IFNγ and TNFα required to prime NLRP3 in keratinocytes. Using two purified SAgs, SEA and SEB, we confirmed that SAgs are indeed highly potent T cell mitogens, and are capable of inducing massive IFNγ and TNFα secretion from donor-derived PBMCs to a similar extent as CD3/CD28 stimulation (Figure S9A). We next performed immunostaining of T cell activation marker CD69 and intracellular IFNγ, as well as ELISA of the conditioned media to quantify the superantigenic activity of all 7 S. aureus strains in our collection (Figure 4A, Figure S9B-C, Figure S10A-D). In agreement with previous epidemiological surveys, SAg activity differs widely among the 7 strains. Three strains, i.e. USA300, 0827 and ATCC29213 were able to activate >3% CD3^+^ T cells, as measured by positive CD69 and IFNγ staining (Figure 4B-C) and induced high levels of IFNγ and TNFα secretion from total PBMCs (Figure S9B-C). These strains are designated as SAg^high^ (Figure 4B-C). 3 other strains (SAg^low^) caused 0.02-0.1% T cell activation (measured by CD69^+^IFNγ^+^), which correlated with low or absent IFNγ and TNFα secretion in PBMCs measured by ELISA (Figure 4B-C, S9B-C). Strain 0360 and *S. epidermidis* (SAg^-ve^) did not cause any T cell activation or induce cytokine secretion. Notably, the MRSA strain USA300 is the only strain that has high SAg activity as well as Hla expression (Figure 4B-C, S9B-C).

**Figure 4.**
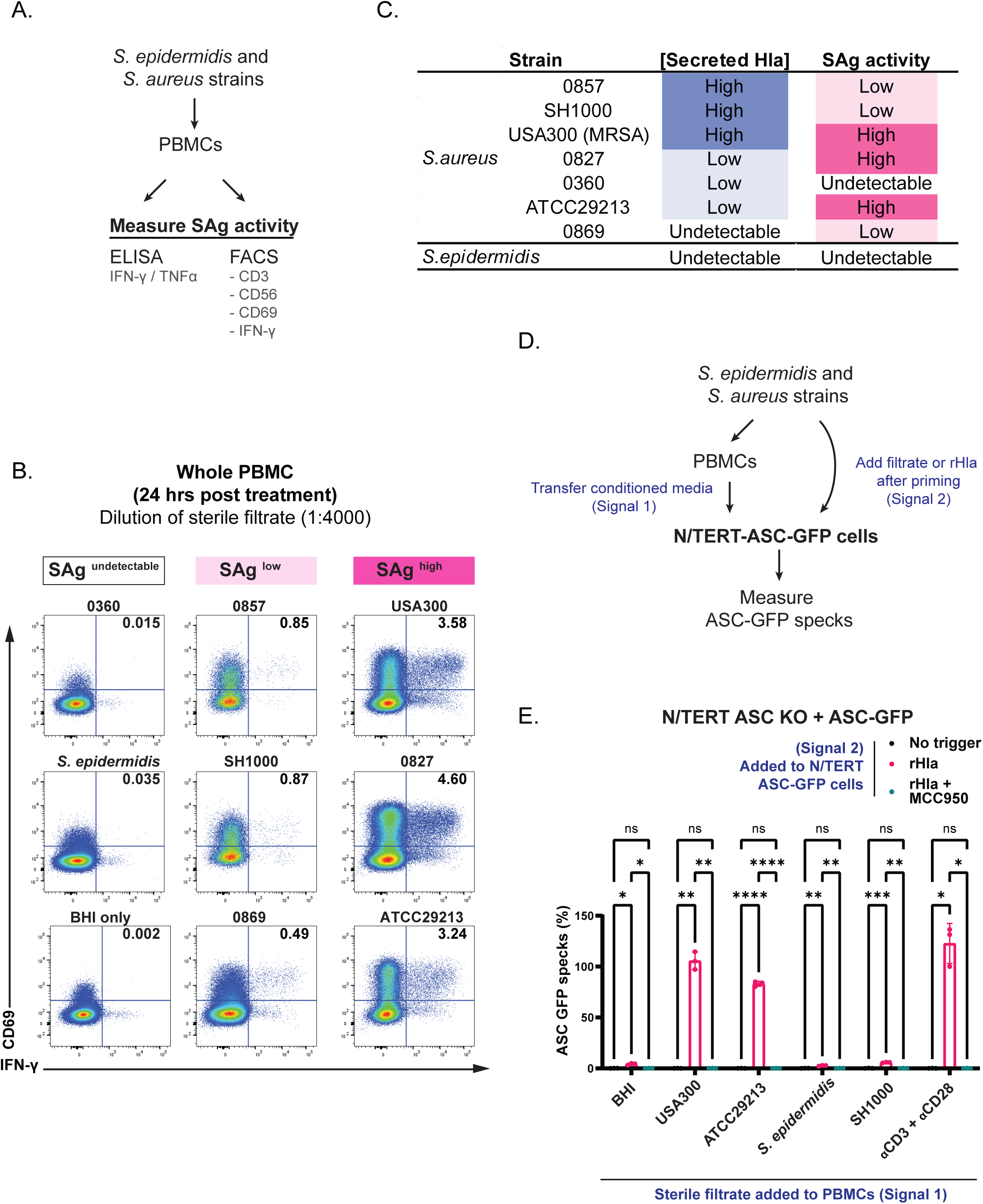
Paracrine IFNγ and TNFα from SAg-activated T cells are a likely source of priming signal for keratinocyte-intrinsic NLRP3. A. Experimental workflow on quantifying the SAg activity of various *S. aureus* strains B. Flow cytometry plots showing the frequency of CD69^+^ and intracellular IFNγ^+^ events among PBMCs stimulated with either anti-CD3/anti-CD28, BHI only (1:4000) or various bacterial sterile filtrates (1:4000) for 24 hrs. Bacterial strains were categorized as SAg^high^ (> 3% CD3^+^ T cells), SAg^low^ (0.02 % - 0.1 % CD3^+^ T cells) and SAg^undetectable^ (und CD3^+^ T cells) C. Summary Hla levels and SAg activities of various bacterial strains. D. Experimental setup of the ‘media transfer’ experiment, assessing the ability of bacterial filtrate treated PBMC conditioned media to prime NLRP3 in keratinocytes E. Results of the media transfer experiment in D, showing the percentage of ASC-GFP specks in N/TERT ASC KO + ASC-GFP cells stimulated overnight in KSFM containing 10% PBMC conditioned media followed by 3 hrs treatment with rHla (1 μg/ml) +/- MCC950 (5 μM) all in the presence of emricasan (5 μM).

We hypothesized that SAg^high^, but not SAg^low^ or SAg^undetectable^ *S. aureus* strains are capable of priming keratinocyte-intrinsic NLRP3 indirectly via PBMC-derived IFNγ and TNFα. A two step ‘media transfer’ model was set up: freshly isolated PBMCs were first stimulated with filtrates of S. aureus cultured differing in SAg activity. The conditioned media were then added to N/TERT ASC-GFP reporter cells for 24 to allow for NLRP3 priming. In the second step, the primed reporter cells were stimulated with either recombinant Hla (Figure 4D). When added to PBMCs (Sigal I, priming step), only SAg^+ve^ strains (e.g. USA300, ATCC29213) enabled the recipient N/TERT-ASC-GFP cells to form ASC-GFP specks in response to recombinant Hla (Signal II) (Figure 4E). These results provide in vitro evidence that SAg-activated T cells are able to provide a sufficient priming signal for the NLRP3 inflammasome in keratinocytes, rendering them susceptible to pyroptosis caused by Hla.

*S. aureus* is one of the most abundant species found in chronic wounds such as atopic dermatitis lesions, diabetic foot ulcers, pressure ulcers and recurrent skin blisters associated with epidermolysis bullosa (EB) (70–72). Many studies have shown that the presence of *S. aureus* virulence factors, including Hla and SAgs are associated with the severity of skin damage and delay in healing (70). Aside from co-occurrence with other bacterial species, recent studies have also found that certain types of wounds, such as those found in EB patients and Buruli ulcers are frequently colonized with multiple *S. aureus* strains (73–75). Our in vitro studies imply that in such scenarios, distinct *S. aureus* strains might cooperate to cause keratinocyte pyroptosis via the combined action of secreted SAg and Hla, which need not be carried by the same strain. To test this hypothesis, we performed the same media transfer experiment where different S. aureus strains were used for both Signal 1 (added to PBMCs) and Signal 2 (added to primed N/TERT-ASC-GFP cells). We observed that SAg^high^ strains such as ATCC29213-treated PBMC media enabled a different strain SH1000 to activate NLRP3 in primed N/TERT cells (Figure 5A, Figure S11A), while neither ATCC29213 or SH1000 was able to prime and activate NLRP3 when used individually as both Signal 1 and Signal 2 (Figure S11B-C). Interestingly, the only strain that was capable to effectively prime and activate NLRP3 was the MRSA strain USA300 (SAg^high^Hla^high^) (Figure S11B-C).

**Figure 5.**
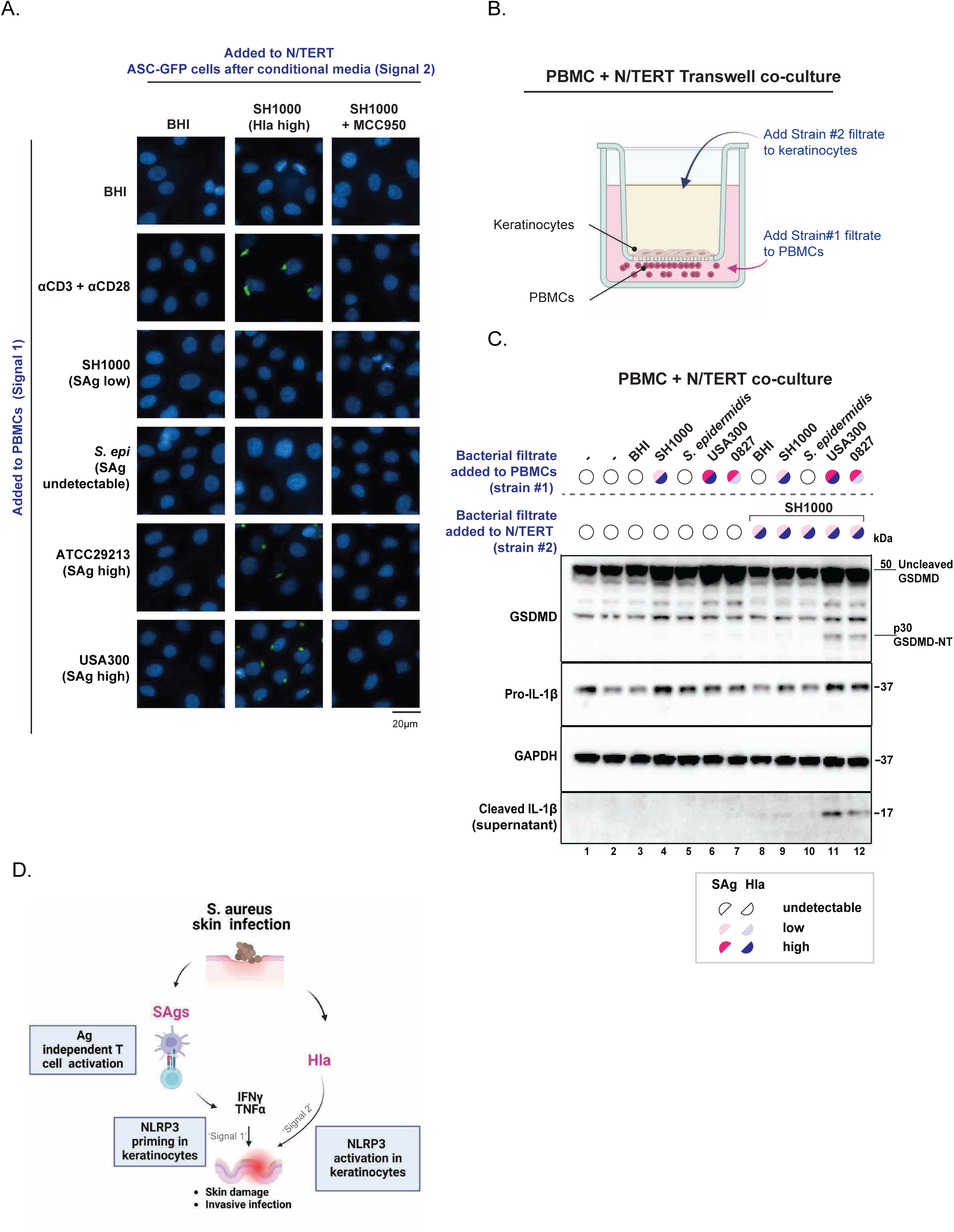
Cooperative activation of the NLRP3 inflammasome by SAg and Hla from diverse *S. aureus* strains. A. Representative images from the media transfer experiment where different strains were used for Signal 1 (added to PBMCs) and Signal 2 (added to N/TERT-ASC-GFP cells) B. Design of the Transwell co-culture assay to study interactions between PBMCs, keratinocytes, and multiple *S. aureus* strains. The strain added to PBMCs is designated Strain #1. The strain added to keratinocytes is designated Strain #2. C. Results of the Transwell assay. The indicated strains were added to PBMCs (Strain #1) 24 hours before the addition of Strain #2 within the Transwells containing a confluent monolayer of N/TERT cells. The N/TERTs were harvested 24 hours after stimulation by direct lysis on the Transwells. Lysates were analyzed by immunoblotting of the indicated antibodies. Supernatant refers to the media within the Transells and was concentrated 10 times before immunoblotting. Results are from one of 2 independent experiments. D. Schematic illustration of the proposed role of NLRP3 inflammasome activation in keratinocytes during *S. aureus* skin infection.

We further confirmed interstrain cooperation using a more direct Transwell assay (Figure 5B). Similar to the media transfer experiment shown above, stimulating PBMCs with SAg^high^Hla^low^ strain 0827 or SAg^high^Hla^high^ strain USA300 enabled SAg^low^Hla^high^ strain SH1000 to induce NLRP3 driven IL-1β secretion and GSDMD cleavage from co-cultured N/TERT cells(Figure 5C, Figure S12A-B). No cooperation was observed between *S. epidermidis* and SH1000.

In summary, our results provide preliminary in vitro evidence that *S. aureus* can employ two well-known exotoxins to mount a coordinated attack on infected skin, culminating in the demise of keratinocytes via NLRP3 driven pyroptosis (Figure 5D). In the first step, SAg causes polyclonal activation of T cells resulting in IFNγ and TNFα secretion. Acting as ‘Signal 1’, these cytokines prime NLRP3 expression in keratinocytes in a paracrine manner. Hla (Signal 2) then enters the cytosol via ADAM10 and activates NLRP3 in the primed keratinocytes leading to inflammasome-driven pyroptosis. As *S. aureus* strains differ widely in SAg and Hla expression, co-infecting strains, when present in the same niche such as chronic skin wounds, could potentially pool their SAg and Hla together to kill keratinocytes via NLRP3-driven pyroptosis. Future experiments are needed to assess the relevance of this model in clinical samples and/or a humanized mouse model.

## DISCUSSION

Our data show that NLRP3 is robustly induced by IFNγ in human immortalized and primary keratinocytes. This rather straightforward result helps resolve the conflicting data in the field. Unlike previously claimed, NLRP3 is neither constitutively expressed nor is it completely absent in human skin keratinocytes. Our results instead show that NLRP3 is strongly repressed under ‘resting’ conditions and sharply induced with the right cytokine milieu. These data thus position IFNγ primed human keratinocytes as an alternate relevant model for investigating endogenous NLRP3 function. This model may allow the field to address certain questions that are not possible with monocytes, macrophages and other cell types of hematopoietic origin. To illustrate its utility, we demonstrated that K+ efflux alone cannot determine the specificity of NLRP3 and NLRP1 activators, with nigercin capable of activating both and *S. aureus* Hla only activating NLRP3. This experiment would not have been possible with macrophage-like cell lines such as THP-1, due to the lack of a functional NLRP1 inflammasome. Our data should serve as the basis for future studies to address the potential crosstalk between the NLRP1 and NLRP3 inflammasomes.

In addition, we provide preliminary in vitro evidence for the interplay between SAg, Hla, T cells and keratinocytes during *S. aureus* skin infection. We propose a model where SAg-intoxicated T cells secretes sufficient IFNγ and TNF to prime NLRP3 expression in keratinocytes, rendering them susceptible to Hla-driven pyroptosis. This pathway might allow avirulent strains to kill keratinocytes cooperatively. If proven in vivo, this mode of ‘interstrain cooperation’ might contribute to the pathogenesis of chronic nonhealing wounds as seen in diabetic ulcers and epidermolysis bullosa patients (73, 76–78).

While it is tempting to speculate that NLRP3 plays a causal role in inflammatory skin diseases such as psoriasis and atopic dermatitis based on our data, we caution against such generalizations. Notably, patients with gain of function mutations in IFNγ signal transducer STAT1 do not suffer from intraepithelial lesions that define NLRP1 GOF mutations (79, 80), suggesting that they might not experience constitutive keratinocyte pyroptosis. In addition, NLRP3 GOF diseases are effectively treated with IL-1 blockade, rather than IFNγ inhibition (81, 82). Therefore, we believe that the pathway described in this study might only apply to contexts with very high local concentrations of IFNγ, such as SAg^+ve^ *S. aureus* infection. The experiments described in this study should guide future clinical research to identify the skin diseases that might benefit from NLRP3 inhibition.

## LIMITATIONS AND CLARIFICATIONS

1. Our study was not intended to identify novel priming or activating signals for NLRP3, as the effects of IFNγ and Hla on other cell types such as macrophages have been extensively documented. Neither do we claim to be the first to demonstrate a role of the NLRP3 inflammasome in keratinocytes. Instead, we focused specifically on known priming and activating molecules in order to resolve the contradictory literature regarding the existence of a functional NLRP3 inflammasome in human (not murine) keratinocytes (Figure 1A). It remains possible that human skin-intrinsic NLRP3 has evolved the capacity to detect entirely novel pathogen-derived molecules originating from skin-tropic microbes that remain to be identified.
2. We did not profile a sufficient number of primary human cell types in our analysis. It is possible that other epithelial cell types, such as those in the lower airway might also upregulate NLRP3 in response to exogenous IFNγ. It would be interesting to test this hypothesis in the context of respiratory bacterial and viral infections.
3. Our study also did not address the mechanistic basis of how NLRP3 is specifically primed by IFNγ in keratinocytes, which likely involves unique epigenetic regulation of the NLRP3 promoter/enhancer region in keratinocytes.
4. A major limitation of our study is the lack of in vivo evidence of the proposed interplay between SAg, Hla, T cells and keratinocyte intrinsic NLRP3. To obtain such evidence, key technical hurdles need to be overcome by the field at large. First, murine skin does not encode inflammasome effector proteins such as ASC, CASP1 or GSDMD (61). In addition, certain *S. aureus* SAgs do not effectively crosslink murine HLA-II (67), at least the HLA allotypes in common mouse strains. A fully humanized mouse model with simultaneous transgenic expression of ASC, CASP1, GSDMD,

NLRP3 in keratinocytes and human HLA might be required to test the results here in vivo. Alternatively, it would be informative to carry out the equivalent of the Transwell co-culture experiments in next-generation human skin organoids with a fully reconstituted vasculature and immune compartment.

## ACKNOWLEDGEMENTS

We thank Prof. Kevin Pethe for extremely helpful advice regarding *S. aureus*, and Prof. Eicke Latz, Prof. Etienne Meunier and Prof. Veit Hornung for advice and guidance on Hla and NLRP3 biology. We would like to thank Dr. Esther Koh (NOBIC, NTU) for assistance with image acquisition and analysis. We also thank Dr. Viduthalai Rasheedkhan Regina (A*STAR, Singapore) and Dr. Morten Kjos (University of Life Sciences, Norway) for providing us with the *S.aureus* mutant strains. The Zhong Lab acknowledges the following funding bodies: Ministry of Education (MOE), Singapore under its Academic Research Fund (AcRF) (MOE RT23/23 and MOE-T2EP30222-0008), National Medical Research Council, Singapore (NMRC 023718-00001), National Research Foundation, Singapore (NRF NRF-NRFF11-2019-0006). The Verma lab acknowledges funding support form the MOE Singapore AcRF Tier 1 (MOE RG94/22) and the Agency for Science, Technology and Research (A*STAR), Skin Research Institute Singapore Grant (SRIS_JRG_1011). J.C. is funded by A*STAR Biomedical Research Council (BMRC) central research funds. J.C. and S.W. are funded by the Asian Skin Microbiome Programme 2.0 IAF-PP grant (H22J1a0040), Y.S.L is supported by the Skin Research Institute Singapore Grant (SRIS_JRG_2010) and A*STAR Graduate Academy (A*GA), National Science Scholarship (NSS).The Robinson Lab acknowledges funding support from the Royal Society (RG\R1\241058) and the Academy of Medical Sciences (SBF009\1153).

---

**Supplementary Figure 1.** A. Immunoblots of NLRP3 and GAPDH (loading control) of WT N/TERT stimulated with various TLR ligands for 16 hrs. Pam3CSK4 (10 ng/ml), HKLM (10^7^ cells/ml), Poly (I:C) high molecular weight (1 μg/ml), LPS (1 μg/ml), Flagellin (1 μg/ml), FSL-1 (50 ng/ml), Imiquimod (1 μg/ml), ssRNA40 (1 μg/ml) and ODN2006 (2.5 μM). B. Immunoblots of NLRP3, MX-1, TrpRS and GAPDH of N/TERTs stimulated with type I IFN (IFNβ), type II IFN (IFNγ) or type III IFN (IFNλ1 or IFNλ3). All IFN stimulation was done at 50 ng/ml +/-TNFα (20 ng/ml) for 16 hrs. C. Immunoblots of NLRP3, GSDMD (full length), pro-caspase-1, ASC, pro-IL-1β, pro-IL-18 and GAPDH in WT N/TERT stimulated for 16 hrs with various cytokines at 50 ng/ml. D. Immunoblots of NLRP3, TrpRs and GAPDH of WT N/TERT stimulated for 16 hrs with IFNy (50 ng/ml) with tofacitinib or deucravacitinib. E. Immunoblots of NLRP1, NLRP3 and GAPDH in WT N/TERT stimulated with IFNγ (50 ng/ml) or TNFα (20 ng/ml) for 16 hrs. Immunoblots show lysates from one of two replicates.

**Supplementary Figure 2.** A. RNAscope staining of paired lesional and nonlesion skin biopsies from a second patient. B. Immunoblots of NLRP3, TrpRs or Ponceau of PBMC or WT N/TERT cells 16 hrs post stimulation with LPS (0.1 μg/ml or 1 μg/ml), IFNγ (50 ng/ml or 200 ng/ml), +/- TNFα (20 ng/ml) or sterile filtrate from SH1000 (1:1000). C. Immunoblots of NLRP3, TrpRs or ponceau (loading control) of kidney organoids and WT N/TERT 16 hrs post stimulation with IFNγ (50 ng/ml) +/- TNFα (20 ng/ml). D. Immunoblots of NLRP3, TrpRs or Ponceau of nasal epithelial cells and WT N/TERT 16 hrs post stimulation with IFNγ (50 ng/ml) + TNFα (20 ng/ml) or LPS (0.1 μg/ml) Immunoblots show lysates from one independent experiment with 3 technical replicates (A and B) or two independent experiments (C) .

**Supplementary Figure 3.** A. IL-1β ELISA of Cas9 control and NLRP3 KO N/TERT cells treated with the indicated cytokines and inflammasome activators. ANS: 1µM. VbP, 10µM. Nigericin: 5µM. B. IL-1β ELISA of Cas9 control NLRP1 and NLRP3 KO cells treated with nigericin with and without MCC950. All cells were primed with IFNγ (50 ng/ml) + TNFα (20 ng/ml). C. Immunoblot of NLRP3 and inflammasome activation markers GSDMD, IL-1β and IL-18 in N/TERT cells treated with the indicated compounds/cytokines. Compound 6p: 1 µM. Nigericin: 5 µM. MCC950: 5 µM

**Supplementary Figure 4.** A. Immunoblot of NLRP3, GSDMD (full length + cleaved), pro-IL-1β, cleaved IL-1β and GAPDH (loading control) of WT N/TERT +/- overnight stimulation with IFNγ (50 ng/ml) + TNFα (20 ng/ml) followed by overnight activation with ANS (1 μM), VbP (3 μM) or rHla (1 μg/ml) +/- MCC950 (5 μM). B. Quantification of ASC-GFP specks in N/TERT ASC KO + ASC-GFP reporter cells upon overnight priming with IFNγ (50 ng/ml) + TNFα (20 ng/ml) followed by 3 hrs activation (1 μM) or rHla (1 μg/ml) +/- MCC950 (5 μM) in the presence of emricasan (5 μM). C. Representative images from B. D. Quantification of ASC-GFP specks in N/TERT ASC KO + ASC-GFP reporter cells upon overnight stimulation with IFNγ (50 ng/ml) + TNFα (20 ng/ml) followed by 3 hrs activation with Vbp (3 μM) or rHla (1 μg/ml) +/- extracellular KCl in the presence of emricasan (5 μM). Error bars represent SEM from three biological replicates, where one replicate refers to independent seeding and treatment of cells. Significance values were calculated based on one-way ANOVA followed by Dunnett’s test. ns, non-significant; **P < 0.01; ****P < 0.0001.

**Supplementary Figure 5.** A. Immunoblots of NLRP3, GSDMD (full length and cleaved), pro-IL-1β, cleaved IL-1β and GAPDH (loading control) of human primary keratinocytes stimulated for 8 hrs with IFNγ (50 ng/ml) followed by overnight activation with CL097 (20 μg/ml) or ANS (1 μM) +/- MCC950 (5 μM), belnacasan (5 μM) or emricasan (5 μM). B. IL-1β ELISA of human primary keratinocytes stimulated for 8 hrs with IFNγ (50 ng/ml) followed by overnight treatment with CL097 (20 μg/ml) or ANS (1 μM) +/- MCC950 (5 μM). Supernatant was harvested 24 hrs post-treatment. C. IL-1β ELISA of human primary keratinocytes stimulated with Hla and the indicated inhibitors. Error bars represent SEM from three biological replicates, where one replicate refers to independent seeding and treatment of cells. Significance values were calculated based on one-way ANOVA followed by Dunnett’s test. ns, non-significant; *P < 0.05; ** P<0.01, *** P<0.001, ****P < 0.0001.

**Supplementary Figure 6.** A. Immunoblots of unprimed unmodified (WT), electroporation control or NEK7 KO N/TERT stimulated overnight with ANS (1 μM), VbP (3 μM) or rHla (1 μg/ml). B. Same as (A), except cells were primed with IFNγ and TNF before stimulation. Immunoblot shows lysates from one experiment performed three independent times.

**Supplementary Figure 7.** A. Growth curves of *S. epidermidis* and various *S. aureus* strains in BHI media as measured by optical density (OD600 nm) across 8 hrs. B. Standard curve correlating optical density with colony-forming units per millilitre (CFU/mL) of the various strains. C. Immunoblot of Hla expression in sterile filtrates obtained from *S. epidermidis* and various *S. aureus* strains. The corresponding CFU/mL for each strain is indicated above the respective blots. D. Immunoblot of Hla expression in sterile filtrates at stated timepoints obtained from mutant *S.aureus* strains from the Nebraska Transposon Mutant Library Project-JE2 USA300 and NE1354. The corresponding CFU/mL for each strain is indicated above the respective blots. E. Representative microscopy images showing DAPI and ASC-GFP specks of N/TERT ASC KO + ASC-GFP stimulated overnight with IFNγ (50 ng/ml) + TNFα (20 ng/ml) followed by activation with sterile filtrates from BHI only, 0857 Hla^high^, SH1000 Hla^high^ or 0827 Hla^low^ for 3 hrs in the presence of emricasan (5 μM). All filtrates were used at a dilution of 1:500. Images are shown from one experiment and are representative of n = 3 independent experiments; scale bar, 20 μm. F. Percentage of ASC-GFP specks forming cells in N/TERT ASC KO + ASC-GFP cells stimulated overnight with IFNγ (50 ng/ml) + TNFα (20 ng/ml) followed by 3 hrs activation with various bacterial sterile filtrates in the presence of emricasan (5 μM). All sterile filtrates were used at a dilution of 1:250 except USA300 (1:50) due to lower CFU count. G. Table of exotoxins (hemolysins and Panton-Valentine leukocidin (PVL)) secreted by *S. epidermidis*, ATCC 29213, USA300 and SH1000 based on genome annotations. Immunoblot shows lysates from one experiment performed three independent times. Error bars represent SEM from three biological replicates, where one replicate refers to independent seeding and treatment of cells. p values were calculated based on one-way ANOVA followed by Dunnett’s test (E). ns, non-significant; **P < 0.01; ****P < 0.0001.

**Supplementary Figure 8.** A. IL-1β ELISA of WT N/TERTs stimulated overnight with IFNγ (50 ng/ml) + TNFα (20 ng/ml) and overnight activation with ANS (1 μM) or sterile filtrates of BHI (1:100), USA300 (1:100) or SH1000 (1:1000) +/-MCC950 (5 μM). Supernatant was harvested 24 hrs post-treatment. B. Schematic illustrating the targeted gene editing of ADAM10 locus in N/TERTs. The upper panel shows WT ADAM10, the sgRNA sequence while the lower part represents the modification of ADAM10 resulting in a KO. C. Immunoblot of NLRP3, GSDMD (full length and cleaved), pro-IL-1β and GAPDH (loading control) of WT vs electroporation control vs ADAM10 KO N/TERT following overnight stimulation with IFNγ (50 ng/ml) and overnight activation with VbP (3 μM), rHla (1 μg/ml) or sterile filtrates from SH1000 or 0857. Sterile filtrates were used at a dilution of 1:1000. Immunoblot shows lysates from one experiment performed three independent times. Error bars represent SEM from three biological replicates, where one replicate refers to independent seeding and treatment of cells. p values were calculated based on one-way ANOVA followed by Dunnett’s test (A). ns, non-significant; **P < 0.01; ****P < 0.0001.

**Supplementary Figure 9.** A. IFNγ and TNF ELISA of PBMCs 48 hrs post stimulation with LPS (0.5 μg/ml), rSEA (1 μg/ml) or rSEB (1 μg/ml) or CD3/ CD28 stimulation. B. IFNγ and TNF ELISA of PBMCs 48 hrs post stimulation. Sterile filtrates were used at a dilution of 1:4000 C. Sterile filtrates were used at a dilution of 1:2000 Error bars represent SEM from three biological replicates, where one replicate refers to independent seeding and treatment of cells. P values were calculated based on one-way ANOVA followed by Dunnett’s test. ns, nonsignificant; ****P < 0.0001 or two-way ANOVA followed by Sidak’s test for multiple pairwise comparisons (B-C) ns, non-significant, *P<0.1,**P<0.01, *** P<0.001, **** P<0.0001

**Supplementary Figure 10.** A. Schematic workflow of PBMC stimulation in preparation of flow cytometry (FACS) analysis. B. Percentage of live and dead PBMCs 24 hrs post-stimulation with bacterial sterile filtrates (1:4000). Cell viability was assessed using live/dead fixable blue dye. C. Gating strategy used in FACS analysis D. Flow cytometry plots displaying the frequency of CD69^+^ and intracellular IFNγ^+^ events in PBMCs. The plots compare unstimulated PBMCs with those stimulated for 24 hrs with anti-CD3/anti-CD28 antibodies or rSEB (0.1 μg/ml).

**Supplementary Figure 11.** A. Quantification of the media transfer experiment shown in Figure 5A. The strains used for Signal I (PBMC stimulation) are marked on the horizontal axis and the strains used in Signal II are marked above the graph. B. Results from an independent media transfer experiment using different strains. C. Summary of the ability of each strain to provide signal 1 and signal 2. Error bars represent SEM from three biological replicates, where one replicate refers to independent seeding and treatment of cells. P values were calculated based on two-way ANOVA followed by Sidak’s test for multiple pairwise comparisons (A-C) ns, non-significant, *P<0.1,*** P<0.001, **** P<0.0001

**Supplementary Figure 12.** A. Immunoblot of NLRP3 and GAPDH from the PBMC and N/TERT co-culture system comparing between 0.4 um vs 1 um transwell pore size. PBMCs were stimulated overnight with various bacterial sterile filtrates (1:4000) or with IFNγ (50 ng/ml) + TNFα (20 ng/ml). N/TERT lysates were processed for this immunoblot. B. Immunoblots of GSDMD (full-length and cleaved), pro-IL-1β, cleaved IL-1β, and GAPDH (loading control) from the PBMC and N/TERT co-culture system. PBMCs were stimulated overnight with rSEA (0.1 μg/ml), rSEB (0.1 μg/ml) or anti-CD3/anti-CD28 antibodies and N/TERTs were stimulated +/- IFNγ (50 ng/ml) + TNFα (20 ng/ml). The following day, N/TERTs were treated overnight with +/- SH1000 filtrate (1:1000). N/TERT lysates and supernatant were processed for this immunoblot. Immunoblots show lysates from one experiment performed three independent times.

## MATERIALS AND METHODS

### Cell Culture and Chemicals

Immortalized human keratinocytes (N/TERT) were provided by H. Rheinwald (MTA). Cells were cultured in Keratinocyte Serum Free Media (Gibco, 17005042), supplemented with 294.4 ng/L human recombinant Epidermal Growth Factor (EGF) (Gibco, 10450-013),25 mg/L Bovine Pituitary Extract (Gibco, 13028-014), and 300 μM of CaCl_2_ (Kanto Chemicals, 07058-00). All cell culture experiments involving immortalized cell lines are conducted under Biological Project Number (BPN-72-2021) approved by the Institutional Biosafety Committee (NTU, Singapore). Primary human keratinocytes were isolated from the foreskin of healthy donors and acquired with informed consent through the Asian Skin Biobank (https://www.a-star.edu.sg/sris/technology-platforms/asian-skin-biobank). All experiments performed with primary keratinocytes were carried out with approval from the A*STAR Human Biomedical Research Office (A*STAR Full IRB-2020-209). Human blood samples from healthy volunteers were collected from Health Sciences Authority (HSA) Singapore. Experiments were performed in compliance with institutional guidelines and were approved by the Institutional Review Board of Nanyang Technological University Singapore (IRB-2018-05-034). The 3D kidney organotypic cultures were performed in collaboration with Yun Xia (Nanyang Technological University, Singapore), and the nasal epithelial cell experiments were conducted in collaboration with Mart Lamers(Duke-NUS Medical School, Singapore). Briefly, nasal epithelial basal cell cultures were derived from healthy volunteers and grown in 2D cultures in expansion medium as described previously(83). Experiments on nasal epithelial cells were performed in compliance with institutional guidelines and were approved by the Institutional Review Board of the National University of Singapore (NUS-IRB-2021-706). Written informed consent was obtained prior to sample collection.

### Bacterial strains and culture conditions

The S. aureus strains used in this study are listed in Figure 3B. Most of the strains were clinical isolates generously provided by Peng Huat Eric Yap (Nanyang Technological University, Singapore). *Staphylococcus aureus* mutant strains were generated by the Nebraska Transposon Mutant Library project (NTML-https://ntml.unmc.edu/) (84)and kindly provided by Morten Kjos (Norwegian University of Life Sciences, Norway) and Viduthalai Rasheedkhan Regina (A*STAR Skin Research Labs, Singapore). We used the *hla-*deficient mutant (NE1354), which was generated from the parental *S. aureus* USA300 strain (JE2 USA300). The strains were cultured overnight in Brain Heart Infusion (BHI) broth at 37°C in a shaking incubator set at 230 r.p.m. Bacterial growth was closely monitored by measuring the OD600 every hour for 8 hrs to track growth and correlate it with toxin induction. Concurrently, 1 mL of culture broth was collected each hour, and the sterile filtrate was obtained by filtering the culture broth through a 0.2 µm syringe filter. To enumerate bacterial colony-forming units (CFUs) after 8 hrs and 16 hrs of growth, the cultures were serially diluted 10-fold by sequentially transferring 20 µL into 180 µL of PBS in triplicate wells, achieving dilutions up to 10 . A 5 µL aliquot of each dilution was spotted in triplicate on LB agar plates and incubated overnight at 37°C. The CFUs were then correlated with the levels of Hla produced by the various strains, which were validated and quantified through immunoblotting assays.

### Harvesting of bacterial sterile filtrates

*S. aureus* strains were incubated at 37°C in a shaking incubator with Brain Heart Infusion (BHI) broth. OD600 readings were recorded hourly until the bacteria reached the late log phase or early stationary phase. At this point, the bacterial cultures were centrifuged at 3000 x g for 10 mins to pellet the cells. The supernatant, containing the secreted toxins, was carefully collected and filtered through a 0.2 µm syringe filter to remove any remaining bacterial cells. The sterile filtrates were then concentrated 10x using 3 kDa MWCO Ultra Centrifugal Filters (Amicon).

### RNAScope staining

RNAScope probing for Human NLRP3 (Hs-NLRP3) was conducted on tissue sections of skin samples from psoriatic patients or controls according to the manufacturer’s protocol (ACDBio). Briefly, slides were first baked in a dry oven at 60°C for 1 hour. Slides were then deparaffinized by fresh xylene and 100% ethanol at room temperature before pre-treatment with hydrogen peroxide and target retrieval reagent at 100°C in a steamer. The HybEZ hybridization oven (ACDBio) was then used to incubate the slides at 40°C for subsequent steps. Slides were treated with the protease plus reagent, followed by the respective hybridization probes (Hs-*NLRP3,* ACD Bio #478021) was our target of interest, bacterial *dapb* (ACD Bio #310043) was used as the negative control and Hs-*PPIB* (ACD Bio #313901) was used for the positive control. AMP reagents were then added after probe-hybridization to amplify the signal, followed by signal detection with the fast red reaction dye. The slides were then counterstained with 25% hematoxylin to elucidate tissue the structure. Slides were completely dried on a 60°C hot plate prior to mounting with Ecomount (Biocare Medical). The slides were visualized and images were captured with a Nikon light microscope.

### Cytokine Analysis

Human IL-1β enzyme-linked immunosorbent assay (ELISA) kit (BD, #557953), human IFNy (Thermo Fisher Scientific, #88-7316-88) and human TNFa (Thermo Fisher Scientific, #88-7346-88) were used in accordance with the manufacturer’s protocols to measure secreted cytokines.

### DRAQ7 Inclusion Assay

Keratinocytes of various genotypes were seeded in a black 12-well plate (Cellvis, P12-1.5P) at a density of 0.8 × 10^5^ cells per well. The following day, cells were stimulated with respective chemicals and stained with 0.3µM of DRAQ7 (ab109202) prior overnight imaging on the high-content screening microscope (Perkin Elmer Operetta CLS imaging system, NTU Optical Bio-Imaging Centre in Nanyang Technological University, Singapore). Brightfield and fluorescence images (DRAQ7 channel, 599 nm/644 nm) were captured every 15 minutes over a period of 15 hours. Analysis was performed using Harmony software (Version 6.0). The ratio of DRAQ7-positive cells to the total cell count was calculated based on data from five fields of view per well, with three wells per treatment. Digital phase contrast was utilized to identify cell borders and determine the number of live cells per field. DRAQ7-positive cells were identified through the DRAQ7 channel.

### CRISPR-Cas9 Knockout

The generation of N/TERT NLRP1 KO and NLRP3 KO have been previously described (Robinson et al., 2023). N/TERT ADAM10 KO and N/TERT NEK7 KO were generated using RNP electroporation. The sgRNA target sequences (5’ to 3’) are: ADAM10 sg1 (GATACCTCTCATATTTACAC), ADAM10 sg 2 (AAGTGTCCCTCTTCATTCGT), NEK7 sg1 (ATAGCCCATATCCGGTCGTA) and NEK7 sg2 (ATAGAAAAGAAAATTGGTCG). Briefly, Cas9 protein and single-guide RNA (sgRNA) were pre-assembled into a ribonucleoprotein (RNP) complex. 2 sgRNAs were used for each electroporation. WT N/TERT cells were mixed with the RNP complex in a nucleocuvette and subjected to electroporation with the Lonza 4D-Nucleofector device. Post-electroporation, cells were immediately transferred into appropriate culture conditions. Knockout efficiency was tested by immunoblot. Alternatively, Sanger sequencing of genomic DNA and overall editing efficiency were determined using the Synthego ICE tool (Synthego Performance Analysis, ICE Analysis. 2019. v2.0. Synthego, https://ice.synthego.com/<u>#/</u>).

### Immunoblotting

Cells were lysed directly with 1x Laemmli buffer (100 µl for 1.8 x 10 cells in a 12-well plate) to prepare whole cell lysates for SDS-PAGE. To visualize GSDMD, cells floating in the supernatant (floaters) were separated after collecting the media, and lysed directly with 1x Laemmli buffer (30 µl for floaters from a 12-well plate). Lysates from the floaters and whole cell lysates were combined in a 1:1 ratio (15 µl floater + 15 µl attached cell lysates) for the detection of GSDMD (full-length and cleaved). To detect cleaved IL-1β, the supernatant was harvested, concentrated using filtered centrifugation (Merck, Amicon Ultra, #UFC500396), and then lysed in the appropriate volume of 5x Laemmli buffer post-concentration. All samples were boiled at 95 for 5 minutes before loading. Proteins were separated by SDS-PAGE and subsequently transferred onto 0.45 μm PVDF membranes (Bio-Rad). The membranes were blocked with 3% milk in PBS, shaking for 1 hour at room temperature, and then incubated overnight at 4 with the appropriate primary antibodies. The following day, the membranes were incubated with the corresponding secondary antibodies on a shaker for 1 hour at room temperature. Protein bands were visualized using the Chemidoc Imaging System (Bio-Rad).

### Immunostaining and fluorescence microscopy

N/TERTs were seeded at a density of 0.8 x 10^5^ cells per well in a 8 well chamber slide (ibidi, 80806). The following day, cells were subjected to the designated treatment conditions. After stimulation, cells were fixed using 4% paraformaldehyde for 15 mins at room temperature and subsequently blocked in a blocking buffer (PBS supplemented with 5% FBS and 0.3% Triton-X-100) for 1 hr at room temperature with gentle shaking. Following the blocking step, cells were briefly rinsed with PBS and incubated overnight at 4 with the respective primary antibodies diluted in an antibody dilution buffer (PBS containing 1% BSA and 0.3% Triton-X-100).The next day, cells were washed three times with PBS for 5 mins each, followed by incubation with the corresponding secondary antibody in the antibody dilution buffer for 1 hour at room temperature on a shaker. To visualize endogenous ASC-specks, primary antibody ASC (Adipogen, AL177) was used, followed by incubation with goat anti-rabbit IgG Alexa Fluor 488-conjugated secondary antibody (Invivogen, A-11008). Endogenous caspase-1 activity was detected using the FAM-FLICA caspase-1 assay kit (ImmunoChemistry Technologies, 9122), following the manufacturer’s protocol.After secondary antibody incubation, cells were washed three times with PBS for 5 mins each and then stained with 4’,6-Diamidino-2-Phenylindole, Dihydrochloride (DAPI, Sigma-Aldrich, MBD0015) for 10 mins at room temperature. Confocal images were captured using the Olympus FV3000 confocal microscope (Olympus Life Sciences).

### ASC-GFP specks quantification

N/TERT ASC GFP KO + ASC-GFP reporter cells were seeded at a density of 0.8 × 10^5^ cells per well in a black 12-well plate (Cellvis, P12-1.5P). Upon treatment, cells were fixed with 4% paraformaldehyde for 15 mins at room temperature and stained with DAPI for 10 mins at room temperature. Images of ASC-GFP specks were acquired in 20 random fields in (DAPI, 358 nm/461 nm) and GFP (469 nm/525 nm) channels under 20x magnification using the Perkin Elmer Operetta CLS imaging system. For quantification of ASC specks, the number of DAPI-stained nuclei and ASC specks in the GFP channel was automatically counted using the Perkin Elmer Operetta CLS imaging system.

### K^+^ efflux inhibition

N/TERT ASC KO + ASC-GFP reporter cells were seeded at a density of 0.8 x 10 cells per well in a 12-well plate. The following day, cells were stimulated overnight with IFNγ (50 ng/ml) and TNFα (20 ng/ml). On the subsequent day, cells were pre-treated with emricasan (5 µM) and extracellular KCl (50 mM) for 15 mins, followed by treatment with recombinant Hla (1 µg/ml) or VbP (3 µM) for 3 hrs. After treatment, cells were fixed and stained with DAPI for ASC speck quantification.

### PBMC isolation

Human blood samples were obtained from healthy volunteers through the HSA Singapore. Peripheral blood mononuclear cells (PBMCs) were isolated using Lymphoprep™ according to a previously optimized protocol [Kizhakeyil et al., 2019]. The isolated PBMCs were washed with Phosphate Buffered Saline (PBS, Nacalai Tesque, Inc., Japan) containing 2% Gibco™ Fetal Bovine Serum (FBS). The cells were then resuspended in complete Roswell Park Memorial Institute (RPMI-1640) culture medium (Invitrogen) supplemented with 10% heat-inactivated FBS and 1% penicillin-streptomycin. Cell counting was performed using a haemocytometer, and cell viability was assessed using trypan blue exclusion, which consistently showed viability greater than 90%. All experiments were conducted in accordance with institutional guidelines and approved by the Institutional Review Board of Nanyang Technological University Singapore (IRB-2018-05-034)

### PBMC stimulation and harvest

T cell activation was induced via TCR engagement. Briefly, a 24-well plate was coated with anti-human CD3 monoclonal antibody (1 µg/mL, Cat#16-0037, eBioscience™) diluted in PBS and incubated for 2 hrs at 37°C. After incubation, the wells were quickly rinsed once with PBS, and PBMCs (1 x 10 cells/well) were seeded in complete RPMI medium supplemented with 0.1 µg/mL soluble anti-human CD28 monoclonal antibody (Cat#16-0289, eBioscience™).The PBMCs were then stimulated with bacterial sterile filtrate for 24 hrs (for flow cytometry) or 48 hrs (for ELISA). Following stimulation, a cell-free supernatant was collected for ELISA, and the cells were harvested for flow cytometry analysis.

### Flow cytometry

PBMCs were seeded at 1 X 10^6^ cells per well in a 24 well plate and stimulated as per conditions described in the figure legends for 24 hrs. Protein transport inhibitor (containing Brefeldin A) (BFA, Biolegend) was added at 5µg per well of cell culture 8 hrs prior to the end of the stimulation for intracellular cytokine staining. Single-cell suspensions were then transferred into 5 mL flow cytometry tubes and incubated with Live/Dead Fixable Blue stain (Molecular Probes) in phosphate buffered saline (PBS) for 15 mins at room temperature in the dark. Cells were then incubated with TruStain FcX (anti-CD16/32, Biolegend) for 15 mins in FACS buffer (2% BSA, 0.02% sodium azide in PBS) at room temperature in the dark. Surface staining was performed in FACS buffer on ice using CD3 (clone UCHT1, BUV395, BD Biosciences), CD56 (clone 5.1H11, APC, Biolegend) and CD69 (clone H1.2F3, FITC, Biolegend) in the dark at room temperature. For intracellular cytokine staining, surface-stained cells were first fixed and permeabilized by the intracellular fixation & permeabilization buffer set (eBioscience) for 30 mins at room temperature in the dark. Intracellular staining was performed in a 1X permeabilization buffer using anti-IFN-γ (Clone REA600, Miltenyi Biotec). Appropriate wash steps were implemented between each staining step with respective buffers. The LSRFortessa X-20 (BD Biosciences) was used for sample acquisition and Flowjo (Treestar) was used for analysis.

### RNAseq sample preparation

N/TERT cells were seeded in a 6-well plate and cultured until they reached 80% confluency, followed by overnight priming with IFNγ (50 ng/ml). Total RNA was extracted from each treatment group using the RNeasy Mini Kit (Cat. #74004; Qiagen). The quantity and quality of the RNA samples were assessed using a NanoDrop spectrophotometer (Thermo Fisher Scientific) and further verified by running the RNA on an agarose gel to check for degradation.RNA sequencing was performed at Macrogen Asia on the NovaSeq 6000 platform. Library construction and sequencing were conducted according to standard protocols by Macrogen Asia. Preprocessing and analysis were also managed by Macrogen Asia, where the sequencing reads were mapped to the reference genome (Homo sapiens, GRCh38) using HISAT2. Transcripts were assembled from the aligned reads using StringTie, and expression profiles were quantified as read counts and normalized values, represented as transcripts per kilobase million (TPM) for each sample, taking into account transcript length and coverage depth.

### Hla Neutralization assay

N/TERT cells were seeded at a density of 0.8 x 10 cells per well in a 12-well plate. The following day, cells were stimulated overnight with IFNγ (50 ng/ml) and TNFα (20 ng/ml). On the subsequent day, anti--hemolysin (20 ug/ml) was pre-incubated with BHI or sterile filtrates of 0857 and SH1000 for (15 mins) to allow binding. After incubation, the mixture of toxin +/-antibody was added to the N/TERTs and incubated overnight.

### Media transfer assay

Freshly isolated PBMCs were stimulated with sterile filtrates from various S. aureus strains for 48 hrs. The cell-free supernatant (conditioned media) was then harvested, and IFNγ and TNFα secretion levels were quantified by ELISA. For the media transfer assay, N/TERT ASC-KO + ASC-GFP cells were seeded at a density of 0.8 x 10 cells/well in a black 24-well plate (Cellvis, P24-1.5P). The following day, conditioned media was added to the cells for overnight stimulation, with 10% of the total volume (500 µL KSFM) being conditioned media. After overnight incubation, the cells were pre-treated with emricasan (5 µM) for 15 mins before treatment with either recombinant Hla (1 µg/ml) or sterile filtrate from S. aureus SH1000 (1:1000 dilution). After 3 hrs, cells were fixed and stained with DAPI for ASC speck quantification using the Perkin Elmer Operetta CLS imaging system.

### Transwell assay

N/TERTs were seeded at a density of 0.6 x 10^5^ cells in 500 µL of KSFM medium per well in a 0.4 µm transwell polyester membrane cell culture insert (CLS3460, Merck). The inserts were placed into a flat-bottom 12-well plate containing 1.5 mL of RPMI medium supplemented with 10% FBS and 1% penicillin-streptomycin per well. The following day, the RPMI medium in the transwell inserts was removed and replaced with freshly isolated PBMCs seeded at a density of 1.5 x 10 cells per well in the supplemented RPMI medium. All treatments of N/TERTs were performed in 500 µL of KSFM medium. After stimulation, the media from the transwell inserts was harvested and centrifuged at maximum speed for 1 min. The cell-free supernatant was collected for the detection of cleaved IL-1β, and the floating cells were lysed in 15 µL of 1x Laemmli buffer for the detection of cleaved GSDMD. Cells remaining in the transwell inserts were lysed directly with 50 µL of 1x Laemmli buffer to prepare whole-cell lysates for SDS-PAGE analysis. Additionally, PBMC-free media was harvested and stored for ELISA.

### Antibodies and drugs used in this study

The following antibodies were used in this study: IL1β p17 specific (Cell Signaling Technology, #83186S), pro–IL-1β (R&D Systems, AF-401-NA), IL-18 (Abcam, ab207324), GAPDH (Santa Cruz Biotechnology, #sc-47724), GSDMDC1 (Novus Biologicals, NBP2-33422), GSDME (Abcam, ab215191), NLRP1 (Biolegend, #9F9B12), NLRP3 (Abcam, ab263899), ASC (Adipogen, AG-25B-0006-C100), Caspase-1 (CST, 3866S), TrpRs (Proteintech, 16081-1-AP), MX-1 (CST, #37849). All horseradish peroxidase (HRP)-conjugated secondary antibodies 4 were purchased from Jackson Immunoresearch (goat anti-mouse IgG: 115-035-166; goat anti-rabbit IgG: 111-035-144). The following drugs and chemicals used are: anisomycin (ANS, MCE, #HY18982), talabostat (VbP, MCE, #HY-13233), MCC950 (MCE, HY-12815A), belnacasan (MCE, HY-13205), emricasan (MCE, HY-10396), LPS-B5 ultrapure (Invivogen, tlrl-pb5lps), TNF (R&D Systems, 210-TA), recombinant IFNy (R&D systems; 285-IF-100), Tofacinib (MCE, HY-40354), Deucravacitinib (MCE, HY-117287), Recombinant Hla (MCE, HY-P2967), Recombinant SEA (Cusabio, CSB-EP350473SMR), Recombinant SEB (Cusabio, CSB-EP314886SMR),

### Statistical Analysis

Statistical analyses were performed using Prism 9 (GraphPad Software, Inc.). Otherwise written, data are reported as mean with SEM. The methods for statistical analysis were included in the figure legend.

## References

1. K. C. Barnett, S. Li, K. Liang, J. P.-Y. Ting, A 360° view of the inflammasome: Mechanisms of activation, cell death, and diseases. Cell 186, 2288–2312 (2023).

2. R. C. Coll, K. Schroder, P. Pelegrín, NLRP3 and pyroptosis blockers for treating inflammatory diseases. Trends Pharmacol. Sci. 43, 653–668 (2022).

3. B. R. Sharma, T.-D. Kanneganti, NLRP3 inflammasome in cancer and metabolic diseases. Nat. Immunol. 22, 550–559 (2021).

4. S. M. Man, T.-D. Kanneganti, Regulation of inflammasome activation. Immunol. Rev. 265, 6–21 (2015).

5. J. Fu, K. Schroder, H. Wu, Mechanistic insights from inflammasome structures. Nat. Rev. Immunol. 24, 518–535 (2024).

6. E. Latz, T. S. Xiao, A. Stutz, Activation and regulation of the inflammasomes. Nat. Rev. Immunol. 13, 397–411 (2013).

7. J. R. Janczy, Mechanisms for Activation and Inhibition of Inflammasomes (2014).

8. P. Broz, V. M. Dixit, Inflammasomes: mechanism of assembly, regulation and signalling. Nat. Rev. Immunol. 16, 407–420 (2016).

9. Y. Zhao, F. Shao, Diverse mechanisms for inflammasome sensing of cytosolic bacteria and bacterial virulence. Curr. Opin. Microbiol. 29, 37–42 (2016).

10. N. Lopes Fischer, N. Naseer, S. Shin, I. E. Brodsky, Effector-triggered immunity and pathogen sensing in metazoans. Nat Microbiol 5, 14–26 (2020).

11. O. Dufies, L. Boyer, RhoGTPases and inflammasomes: Guardians of effector-triggered immunity. PLoS Pathog. 17, e1009504 (2021).

12. W. Jing, J. Lo Pilato, C. Kay, S. M. Man, Activation mechanisms of inflammasomes by bacterial toxins. Cell. Microbiol. 23, e13309 (2021).

13. S. Alehashemi, R. Goldbach-Mansky, Human Autoinflammatory Diseases Mediated by NLRP3-, Pyrin-, NLRP1-, and NLRC4-Inflammasome Dysregulation Updates on Diagnosis, Treatment, and the Respective Roles of IL-1 and IL-18. Front. Immunol. 11, 1840 (2020).

14. C. R. Harapas, A. Steiner, S. Davidson, S. L. Masters, An Update on Autoinflammatory Diseases: Inflammasomopathies. Curr. Rheumatol. Rep. 20, 40 (2018).

15. F.-G. Bauernfeind, The NLRP3 Inflammasome: Molecular Mechanisms of Activation and Transcriptional Regulation (2011).

16. S. Le Jan, et al., IL-23/IL-17 axis activates IL-1β-associated inflammasome in macrophages and generates an auto-inflammatory response in a subgroup of patients with bullous Pemphigoid. Front. Immunol. 10, 1972 (2019).

17. A. Wree, et al., NLRP3 inflammasome driven liver injury and fibrosis: Roles of IL-17 and TNF in mice. Hepatology 67, 736–749 (2018).

18. H.-H. Chi, et al., IL-36 signaling facilitates activation of the NLRP3 inflammasome and IL-23/IL-17 axis in renal inflammation and fibrosis. J. Am. Soc. Nephrol. 28, 2022–2037 (2017).

19. M. D. McGeough, et al., TNF regulates transcription of NLRP3 inflammasome components and inflammatory molecules in cryopyrinopathies. J. Clin. Invest. 127, 4488–4497 (2017).

20. D. Fox, et al., Bacillus cereus non-haemolytic enterotoxin activates the NLRP3 inflammasome. Nat. Commun. 11, 760 (2020).

21. W. Jing, et al., Clostridium septicum α-toxin activates the NLRP3 inflammasome by engaging GPI-anchored proteins. Sci. Immunol. 7, eabm1803 (2022).

22. J. Zheng, et al., A novel function of NLRP3 independent of inflammasome as a key transcription factor of IL-33 in epithelial cells of atopic dermatitis. Cell Death Dis. 12, 871 (2021).

23. K. Mähönen, et al., Activation of NLRP3 inflammasome in the skin of patients with systemic and cutaneous lupus erythematosus. Acta Derm. Venereol. 102, adv00708 (2022).

24. S. Li, et al., Activated NLR family pyrin domain containing 3 (NLRP3) inflammasome in keratinocytes promotes cutaneous T-cell response in patients with vitiligo. J. Allergy Clin. Immunol. 145, 632–645 (2020).

25. R. D. Correia-Silva, et al., Regulatory role of annexin A1 in NLRP3 inflammasome activation in atopic dermatitis: insights from keratinocytes in human and murine studies. J. Mol. Med. 103, 435–451 (2025).

26. X. Dai, M. Tohyama, M. Murakami, K. Sayama, Epidermal keratinocytes sense dsRNA via the NLRP3 inflammasome, mediating interleukin (IL)-1β and IL-18 release. Exp. Dermatol. 26, 904–911 (2017).

27. M.-S. Lee, et al., Shiga toxins activate the NLRP3 inflammasome pathway to promote both production of the proinflammatory cytokine interleukin-1β and apoptotic cell death. Infect. Immun. 84, 172–186 (2015).

28. D. A. Bachovchin, NLRP1: a jack of all trades, or a master of one? Mol. Cell 81, 423–425 (2021).

29. C. Y. Taabazuing, A. R. Griswold, D. A. Bachovchin, The NLRP1 and CARD8 inflammasomes. Immunol. Rev. 297, 13–25 (2020).

30. S. Bauernfried, V. Hornung, Human NLRP1: From the shadows to center stage. J. Exp. Med. 219 (2022).

31. S. Grandemange, et al., A new autoinflammatory and autoimmune syndrome associated with NLRP1 mutations: NAIAD (NLRP1-associated autoinflammation with arthritis and dyskeratosis). Ann. Rheum. Dis. 76, 1191–1198 (2017).

32. F. L. Zhong, et al., Germline NLRP1 Mutations Cause Skin Inflammatory and Cancer Susceptibility Syndromes via Inflammasome Activation. Cell 167, 187–202.e17 (2016).

33. S. B. Drutman, et al., Homozygous NLRP1 gain-of-function mutation in siblings with a syndromic form of recurrent respiratory papillomatosis. Proc. Natl. Acad. Sci. U. S. A. 116, 19055–19063 (2019).

34. E. Bostan, O. Gokoz, N. Atakan, The role of NLRP1 and NLRP3 inflammasomes in the etiopathogeneses of pityriasis lichenoides chronica and mycosis fungoides: an immunohistochemical study. Arch. Derm. Res. 315, 231–239 (2023).

35. GTEx Consortium, The Genotype-Tissue Expression (GTEx) project. Nat. Genet. 45, 580–585 (2013).

36. ENCODE Project Consortium, An integrated encyclopedia of DNA elements in the human genome. Nature 489, 57–74 (2012).

37. M. Karlsson, et al., A single-cell type transcriptomics map of human tissues. Sci. Adv. 7, eabh2169 (2021).

38. C. B. Xie, et al., Complement membrane attack complexes assemble NLRP3 inflammasomes triggering IL-1 activation of IFN-γ-primed human endothelium. Circ. Res. 124, 1747–1759 (2019).

39. H. He, et al., Microglial priming by IFN-γ involves STAT1-mediated activation of the NLRP3 inflammasome. CNS Neurosci. Ther. 30, e70061 (2024).

40. A. H. Karaba, et al., Herpes simplex virus type 1 inflammasome activation in proinflammatory human macrophages is dependent on NLRP3, ASC, and caspase-1. PLoS One 15, e0229570 (2020).

41. R. Karki, et al., Synergism of TNF-α and IFN-γ Triggers Inflammatory Cell Death, Tissue Damage, and Mortality in SARS-CoV-2 Infection and Cytokine Shock Syndromes. Cell 184, 149–168.e17 (2021).

42. M. Takahama, et al., A pairwise cytokine code explains the organism-wide response to sepsis. Nat. Immunol. 1–14 (2024).

43. C. Conrad, et al., TNF blockade induces a dysregulated type I interferon response without autoimmunity in paradoxical psoriasis. Nat. Commun. 9, 25 (2018).

44. N. N. Mehta, et al., IFN-γ and TNF-α synergism may provide a link between psoriasis and inflammatory atherogenesis. Sci. Rep. 7, 13831 (2017).

45. N. O. Kurtovic, E. K. Halilovic, Serum concentrations of Interferon gamma (IFN-γ) in patients with psoriasis: Correlation with clinical type and severity of the disease. Med. Arch. 72, 410–413 (2018).

46. P. Rozario, et al., Mechanistic basis for potassium efflux-driven activation of the human NLRP1 inflammasome. Proc. Natl. Acad. Sci. U. S. A. 121, e2309579121 (2024).

47. G. Fenini, et al., The p38 Mitogen-Activated Protein Kinase Critically Regulates Human Keratinocyte Inflammasome Activation. J. Invest. Dermatol. 138, 1380–1390 (2018).

48. R. R. Craven, et al., Staphylococcus aureus alpha-hemolysin activates the NLRP3-inflammasome in human and mouse monocytic cells. PLoS One 4, e7446 (2009).

49. C. Kebaier, et al., Staphylococcus aureus α-Hemolysin Mediates Virulence in a Murine Model of Severe Pneumonia Through Activation of the NLRP3 Inflammasome. J. Infect. Dis. 205, 807–817 (2012).

50. L. Song, et al., Structure of staphylococcal alpha-hemolysin, a heptameric transmembrane pore. Science 274, 1859–1866 (1996).

51. J. Harder, et al., Activation of the Nlrp3 inflammasome by Streptococcus pyogenes requires streptolysin O and NF-kappa B activation but proceeds independently of TLR signaling and P2X7 receptor. J. Immunol. 183, 5823– 5829 (2009).

52. C. J. Groß, et al., K Efflux-Independent NLRP3 Inflammasome Activation by Small Molecules Targeting Mitochondria. Immunity 45, 761–773 (2016).

53. L. K. Billingham, et al., Mitochondrial electron transport chain is necessary for NLRP3 inflammasome activation. Nat. Immunol. 23, 692–704 (2022).

54. Y. He, M. Y. Zeng, D. Yang, B. Motro, G. Núñez, NEK7 is an essential mediator of NLRP3 activation downstream of potassium efflux. Nature 530, 354–357 (2016).

55. H. Shi, et al., NLRP3 activation and mitosis are mutually exclusive events coordinated by NEK7, a new inflammasome component. Nat. Immunol. 17, 250–258 (2016).

56. J. L. Schmid-Burgk, et al., A Genome-wide CRISPR (Clustered Regularly Interspaced Short Palindromic Repeats) Screen Identifies NEK7 as an Essential Component of NLRP3 Inflammasome Activation. J. Biol. Chem. 291, 103–109 (2016).

57. N. A. Schmacke, et al., IKKβ primes inflammasome formation by recruiting NLRP3 to the trans-Golgi network. Immunity 55, 2271–2284.e7 (2022).

58. I. Muela-Zarzuela, et al., NEK7 activates the NLRP1 Inflammasome. *bioRxiv* 2022.12.21.521400 (2022).

59. X. Wang, W. J. Eagen, J. C. Lee, Orchestration of human macrophage NLRP3 inflammasome activation by Staphylococcus aureus extracellular vesicles. Proc. Natl. Acad. Sci. U. S. A. 117, 3174–3184 (2020).

60. D. Holzinger, et al., Staphylococcus aureus Panton-Valentine leukocidin induces an inflammatory response in human phagocytes via the NLRP3 inflammasome. J. Leukoc. Biol. 92, 1069–1081 (2012).

61. J. Sand, et al., Expression of inflammasome proteins and inflammasome activation occurs in human, but not in murine keratinocytes. Cell Death Dis. 9, 24 (2018).

62. G. A. Wilke, J. Bubeck Wardenburg, Role of a disintegrin and metalloprotease 10 in Staphylococcus aureus alpha-hemolysin-mediated cellular injury. Proc. Natl. Acad. Sci. U. S. A. 107, 13473–13478 (2010).

63. A. J. Greaney, S. H. Leppla, M. Moayeri, Bacterial exotoxins and the inflammasome. Front. Immunol. 6, 570 (2015).

64. A. Babbar, Streptococcal Superantigens (Springer, 2015).

65. T. Krakauer, Staphylococcal Superantigens: Pyrogenic Toxins Induce Toxic Shock. Toxins 11 (2019).

66. Staphylococcal Enterotoxins, Toxic Shock Syndrome Toxin-I, and Streptococcal Pyrogenic Exotoxins: Some Basic Biology of Bacterial Superantigens (2003).

67. S. W. Tuffs, et al., Superantigens promote bloodstream infection by eliciting pathogenic interferon-gamma production. Proc. Natl. Acad. Sci. U. S. A. 119 (2022).

68. S. X. Xu, J. K. McCormick, Staphylococcal superantigens in colonization and disease. Front. Cell. Infect. Microbiol. 2, 52 (2012).

69. W. Salgado-Pabón, et al., Superantigens are critical for Staphylococcus aureus Infective endocarditis, sepsis, and acute kidney injury. MBio 4 (2013).

70. L. R. Kalan, et al., Strain-and species-level variation in the microbiome of diabetic wounds is associated with clinical outcomes and therapeutic efficacy. Cell Host Microbe 25, 641–655.e5 (2019).

71. K. Gjødsbøl, et al., Multiple bacterial species reside in chronic wounds: a longitudinal study. Int. Wound J. 3, 225–231 (2006).

72. Multi-Omics Analysis of Wound Microbiome and Staphylococcus aureus in Pressure Ulcer.

73. M. M. van der Kooi-Pol, et al., Topography of distinct Staphylococcus aureus types in chronic wounds of patients with epidermolysis bullosa. PLoS One 8, e67272 (2013).

74. N. A. Amissah, et al., Genetic diversity of Staphylococcus aureus in Buruli ulcer. PLoS Negl. Trop. Dis. 9, e0003421 (2015).

75. A. N. García-Pérez, et al., From the wound to the bench: exoproteome interplay between wound-colonizing Staphylococcus aureus strains and co-existing bacteria. Virulence 9, 363–378 (2018).

76. R. Blakytny, E. Jude, The molecular biology of chronic wounds and delayed healing in diabetes. Diabet. Med. 23, 594–608 (2006).

77. A. Bardhan, et al., Epidermolysis bullosa. Nat. Rev. Dis. Primers 6, 78 (2020).

78. K. Shettigar, T. S. Murali, Virulence factors and clonal diversity of Staphylococcus aureus in colonization and wound infection with emphasis on diabetic foot infection. Eur. J. Clin. Microbiol. Infect. Dis. 39, 2235–2246 (2020).

79. S. Okada, et al., Human STAT1 Gain-of-Function Heterozygous Mutations: Chronic Mucocutaneous Candidiasis and Type I Interferonopathy. J. Clin. Immunol. 40, 1065–1081 (2020).

80. Y. Mizoguchi, S. Okada, Inborn errors of STAT1 immunity. Curr. Opin. Immunol. 72, 59–64 (2021).

81. N. Martinez-Quiles, R. Goldbach-Mansky, Updates on autoinflammatory diseases. Curr. Opin. Immunol. 55, 97–105 (2018).

82. M. S. J. Mangan, et al., Targeting the NLRP3 inflammasome in inflammatory diseases. Nat. Rev. Drug Discov. 17, 688 (2018).

83. G. D. Amatngalim, et al., Measuring cystic fibrosis drug responses in organoids derived from 2D differentiated nasal epithelia. Life Sci Alliance 5 (2022).

84. P. D. Fey, et al., A genetic resource for rapid and comprehensive phenotype screening of nonessential Staphylococcus aureus genes. MBio 4, e00537– 12 (2013).

