## Supplemental Figures S1-S12 for "Clarifying the priming requirement and activator specificity for the NLRP3 inflammasome in human keratinocytes in vitro"

### Fig S1(For Fig 1)

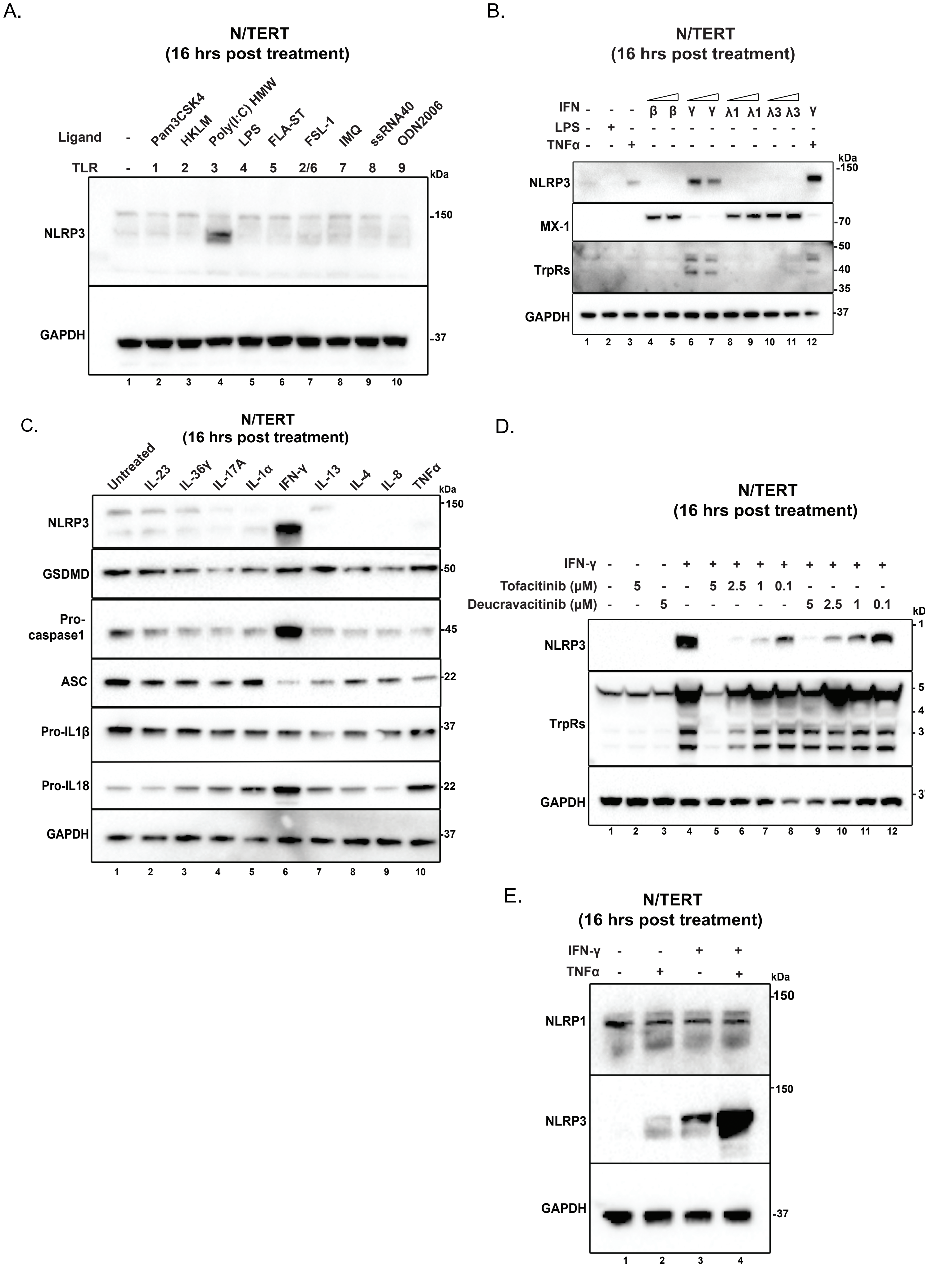

### Fig S2 (For Fig 1)

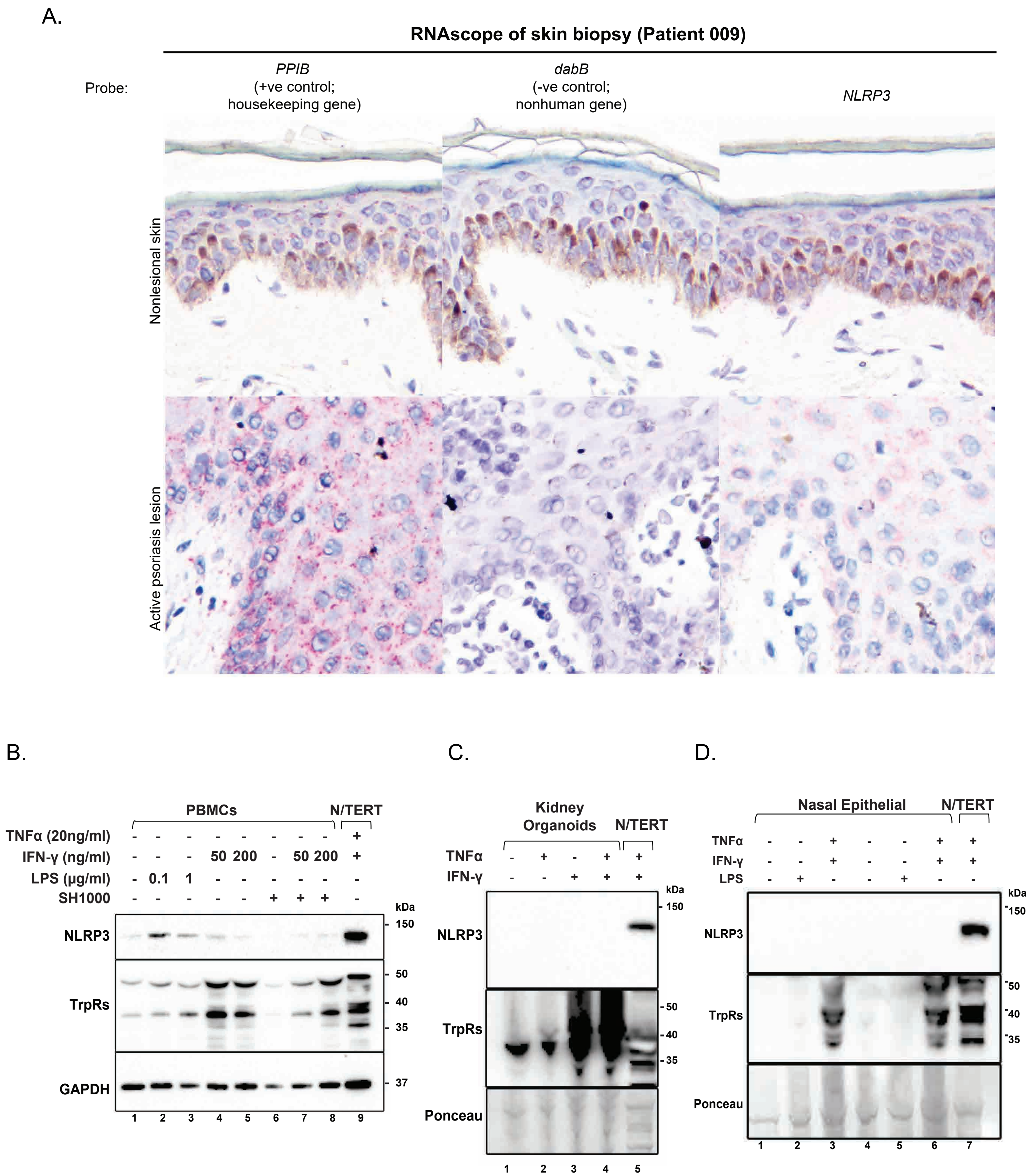

### Fig S3

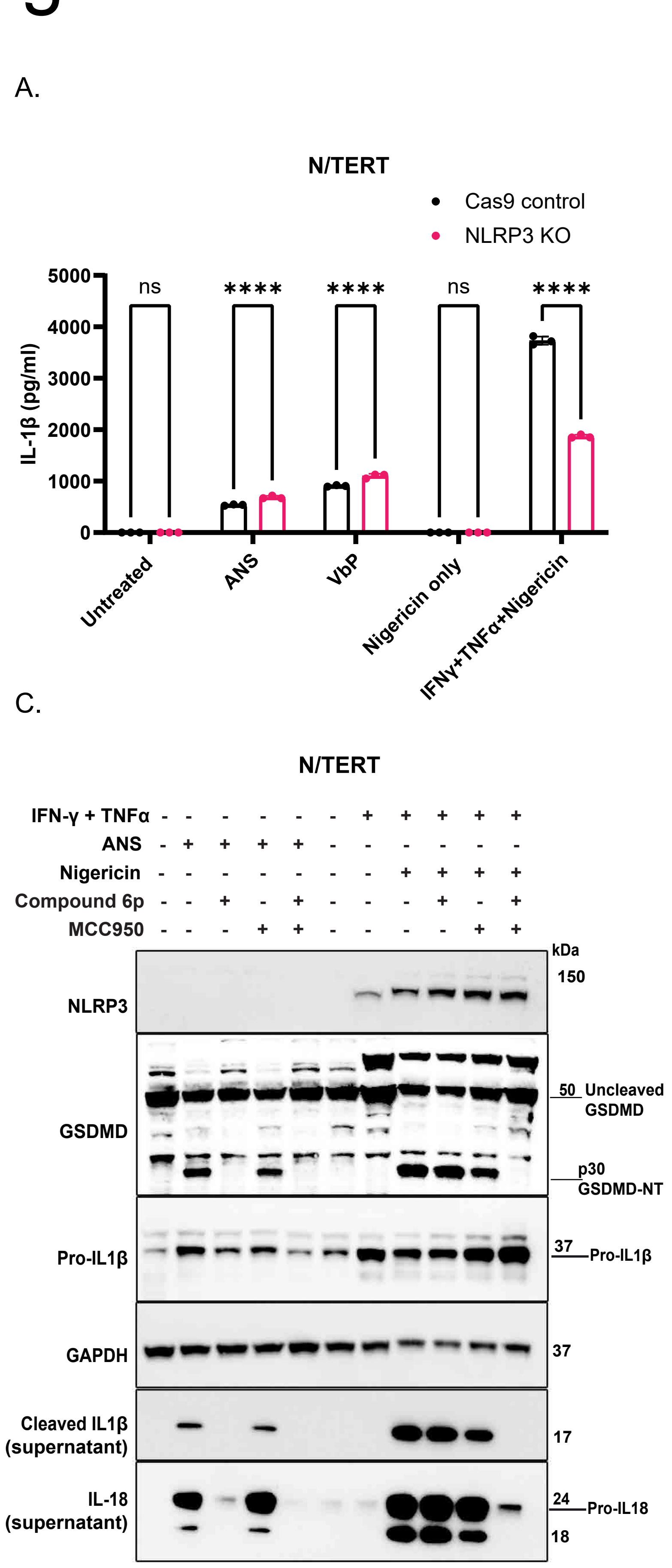

A.
N/TERT
Cas9 control
NLRP3 KO
IL-1β (pg/ml)
5000
4000
3000
2000
1000
0
ns
****
****
ns
****
Untreated
ANS
VbP
Nigericin only
IFNγ+TNFα+Nigericin
B.
N/TERT
(IFN-γ+TNFα primed)
IL-1β (pg/mL)
1000
800
600
400
200
0
ns
ns
ns
ns
ns
ns
****
****
****
***
***
*
Cas9 control
NLRP1 KO
Nigericin
MCC950
C.
N/TERT
IFN-γ + TNFα
ANS
Nigericin
Compound 6p
MCC950
kDa
NLRP3
150
GSDMD
50 Uncleaved GSDMD
p30 GSDMD-NT
Pro-IL1β
37 Pro-IL1β
GAPDH
37
Cleaved IL1β (supernatant)
17
IL-18 (supernatant)
24 Pro-IL18
18

### Fig S4(For Fig 2)

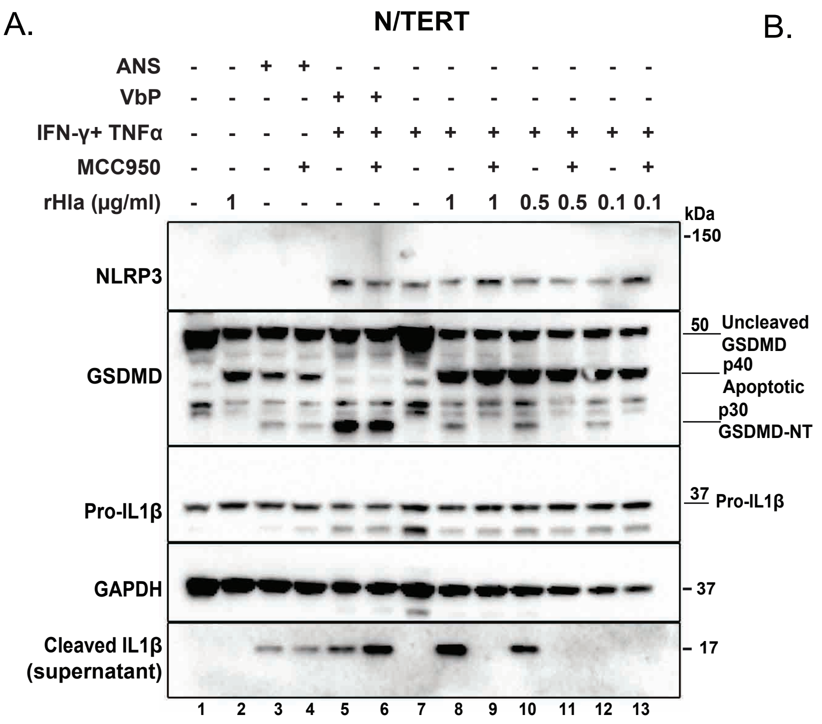


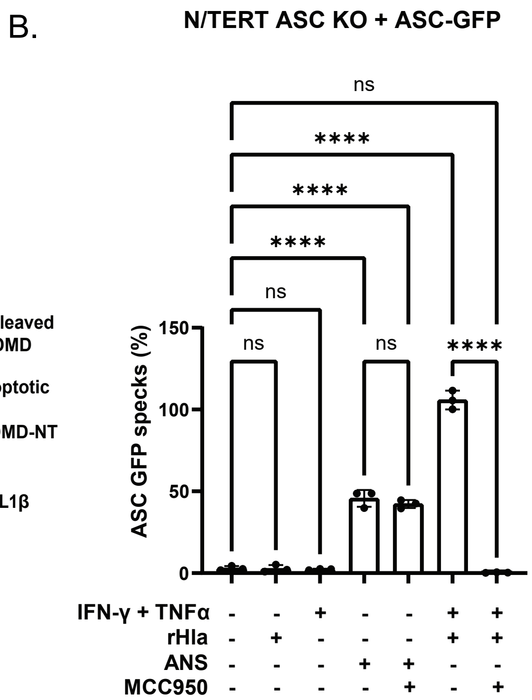


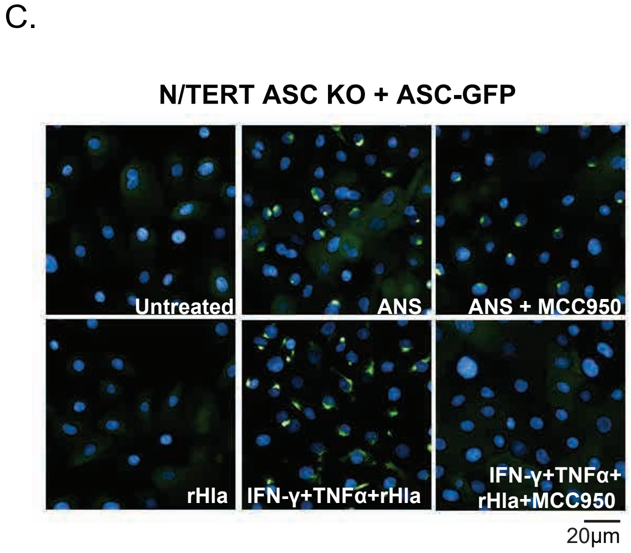


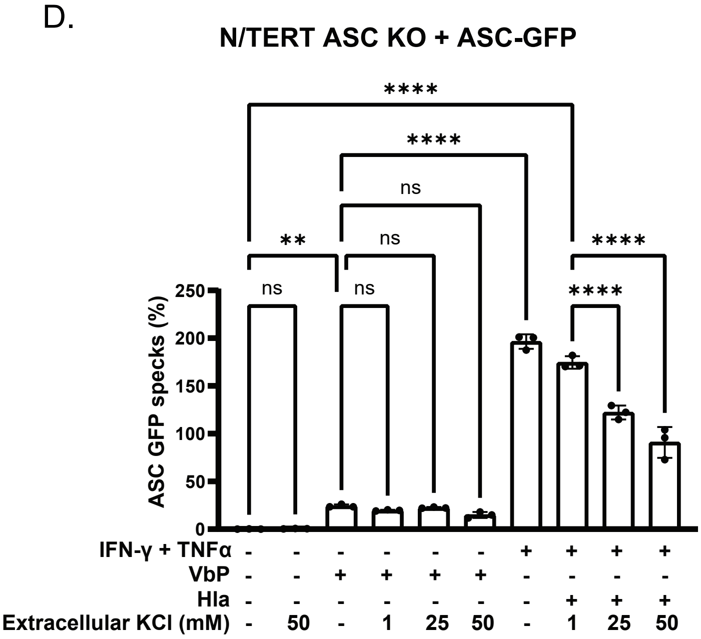

### Fig S5 (For Fig 2)

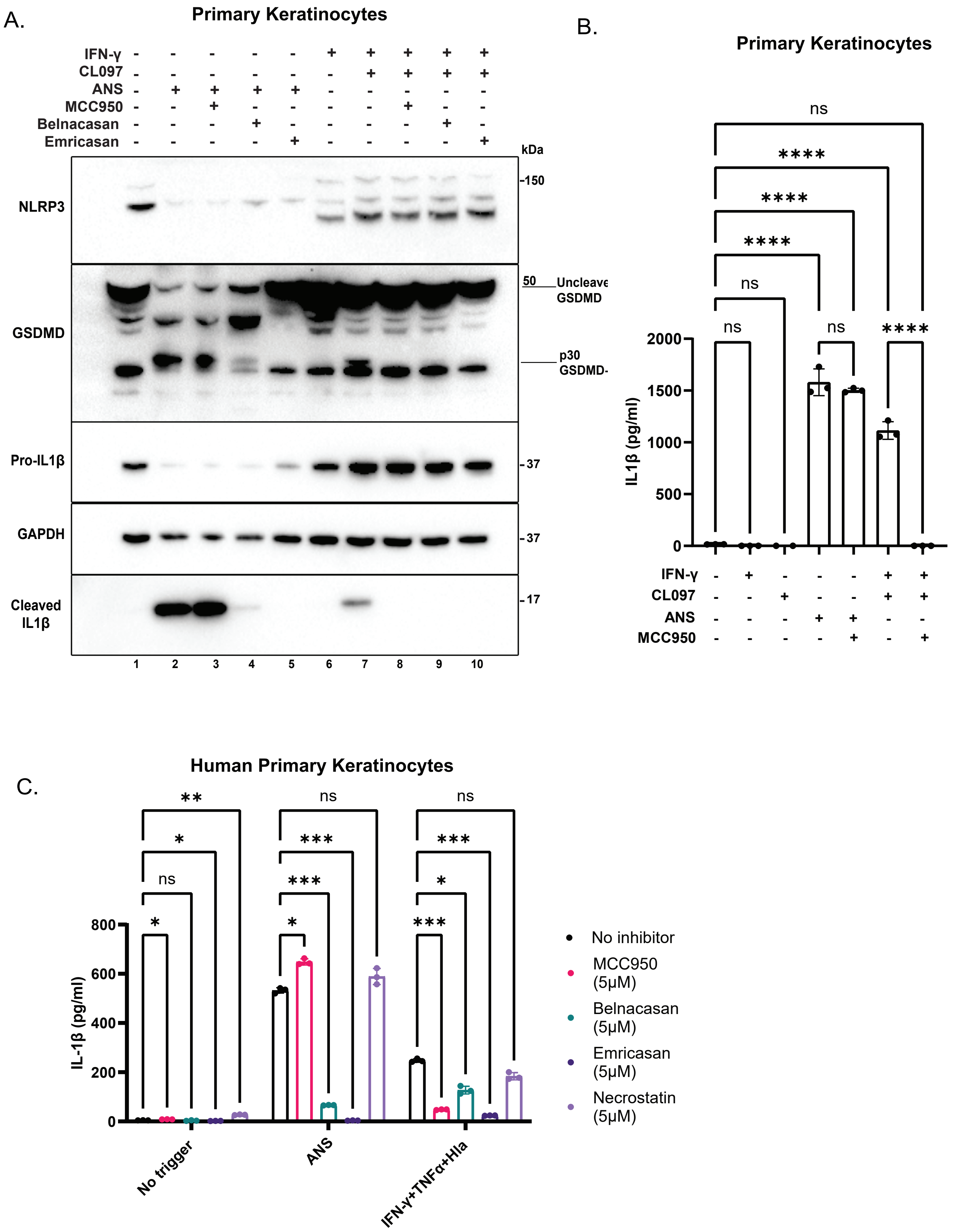

### Fig S6 (For Fig 2)

A.

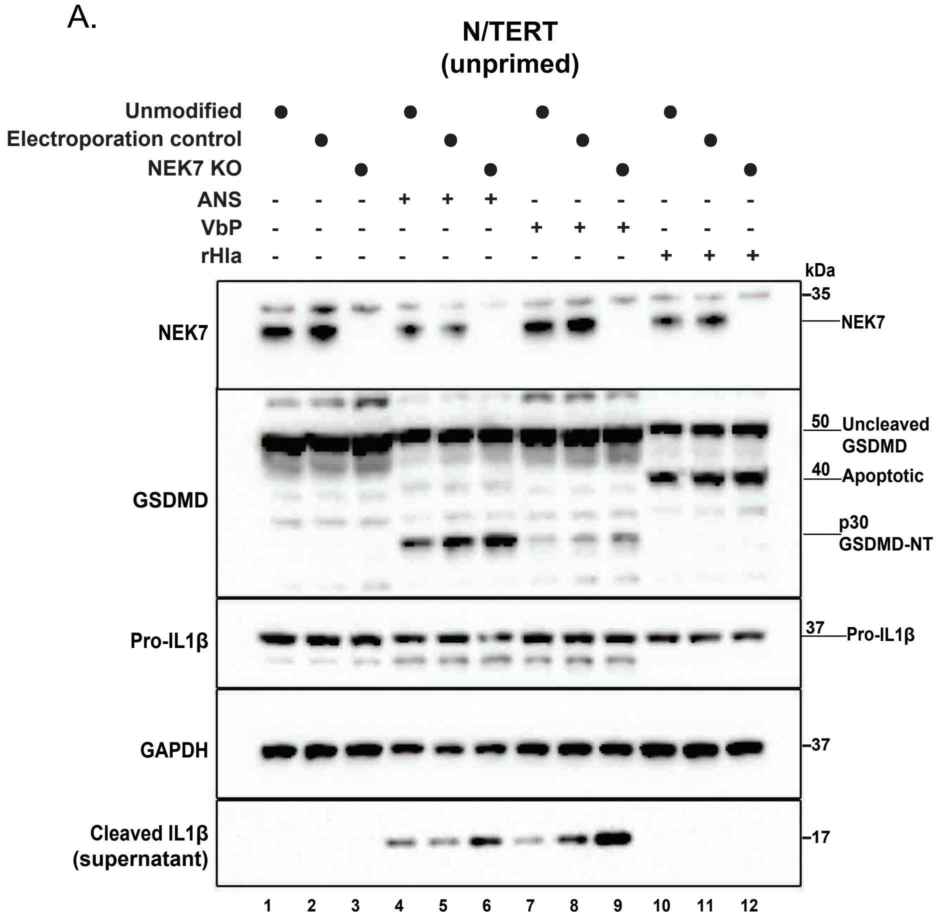


B.

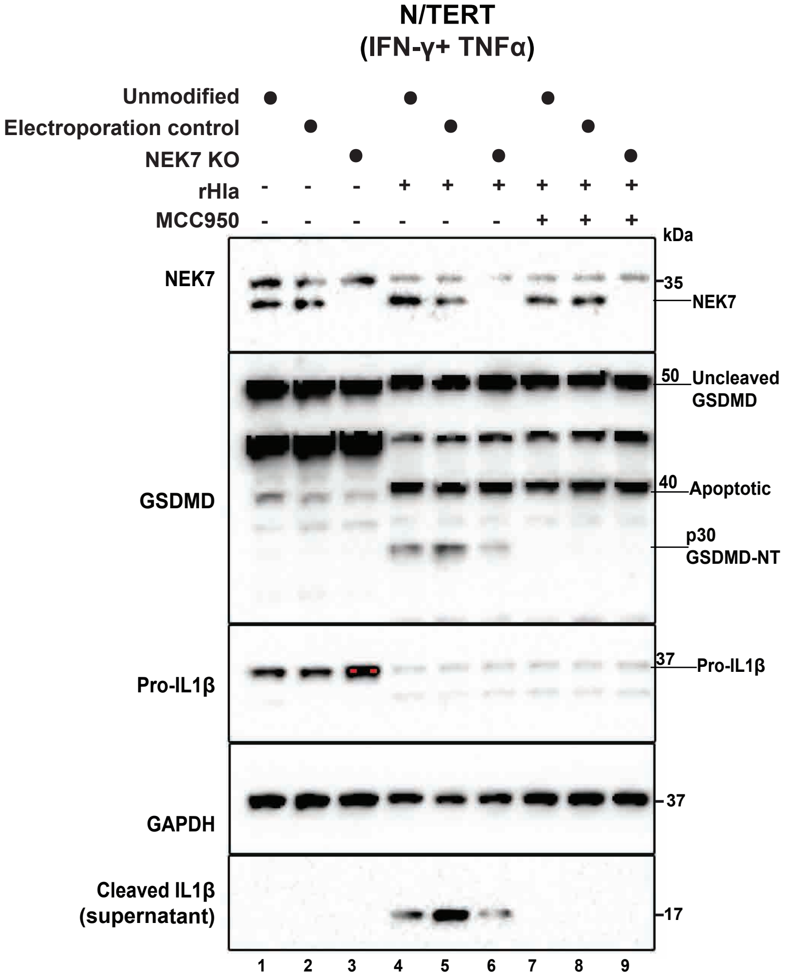

### Supplementary Fig 7 (For Fig 3)

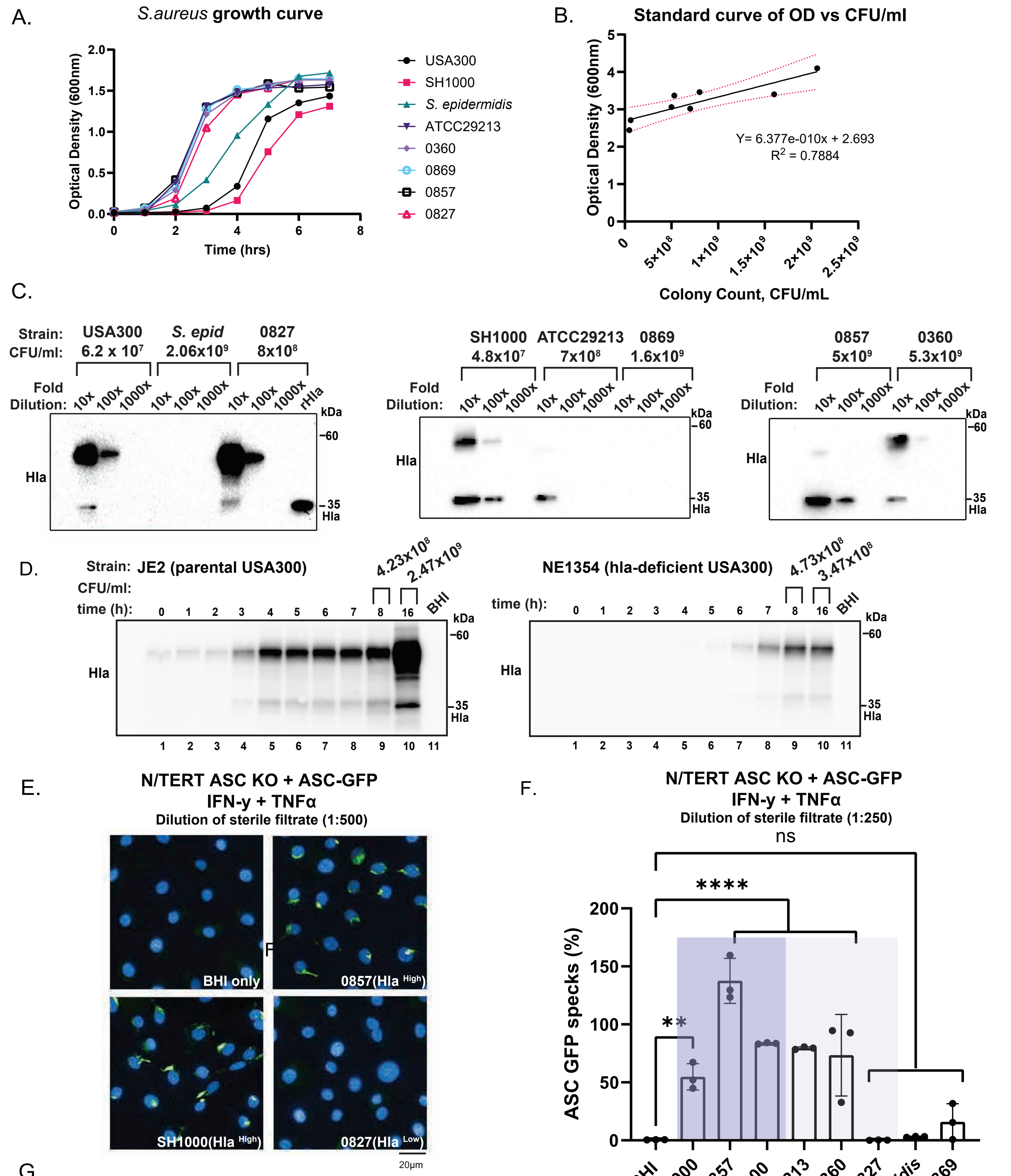


*Sterile filtrate of USA300 was used at 1:50 due to low CFU

G.

**Presence of PFTs based on genome annotation**

| Strain | | Hemolysins | | | Panton–Valentine leukocidin (PVL) | |
|---|---|---|---|---|---|---|
| | | Hla | Hlb | Hly | lukS- PV | lukF- PV |
| *S.aureus* | ATCC29213 | Yes | No | Yes | No | No |
| | USA300 | Yes | Yes | No | Yes | Yes |
| | SH1000 | Yes | No | Yes | No | No |
| *S. epidermidis* | | No | No | No | No | No |

### Fig S8 (For Fig 3)

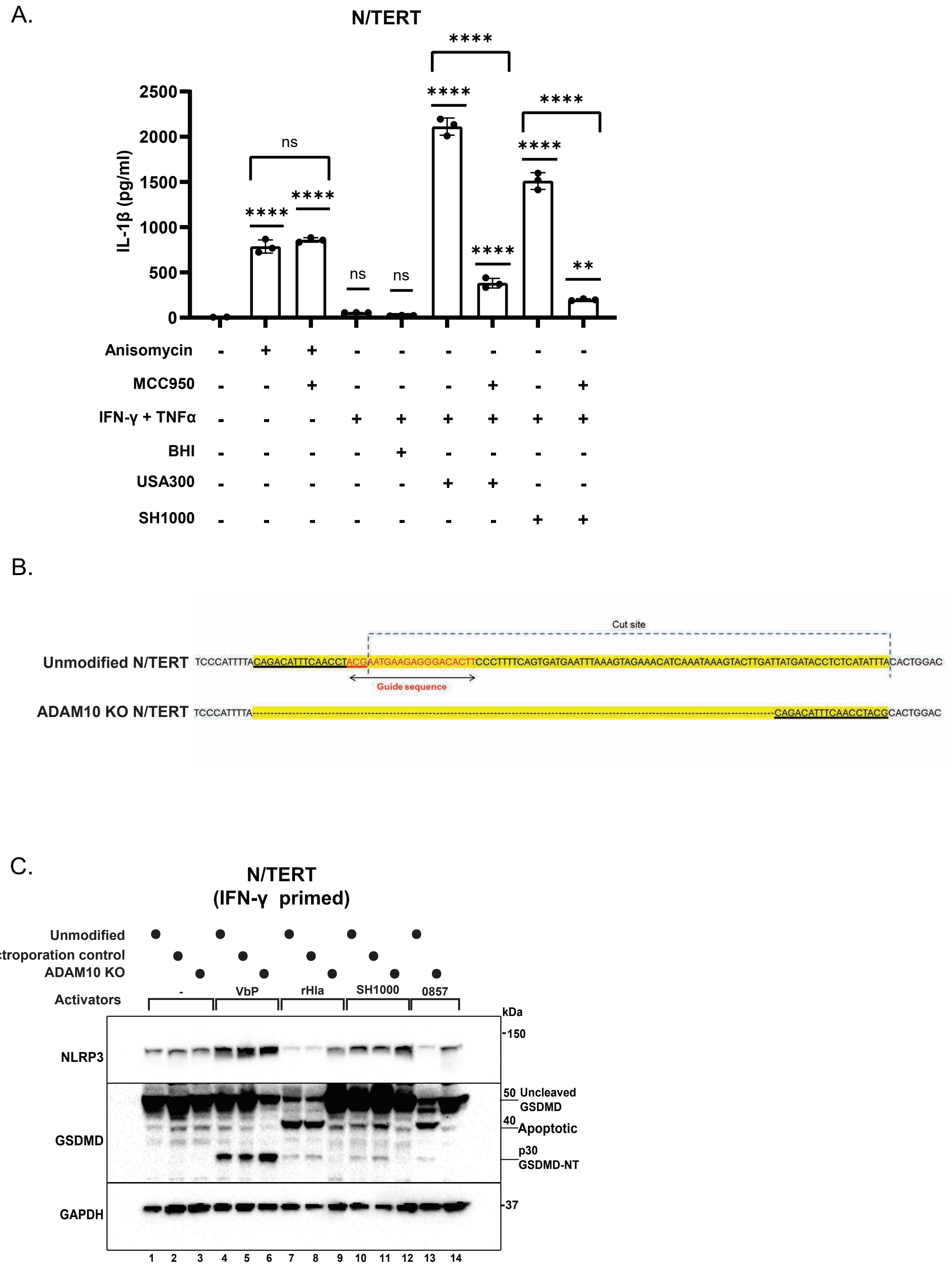

### Fig S9 (For Fig 4)

A.

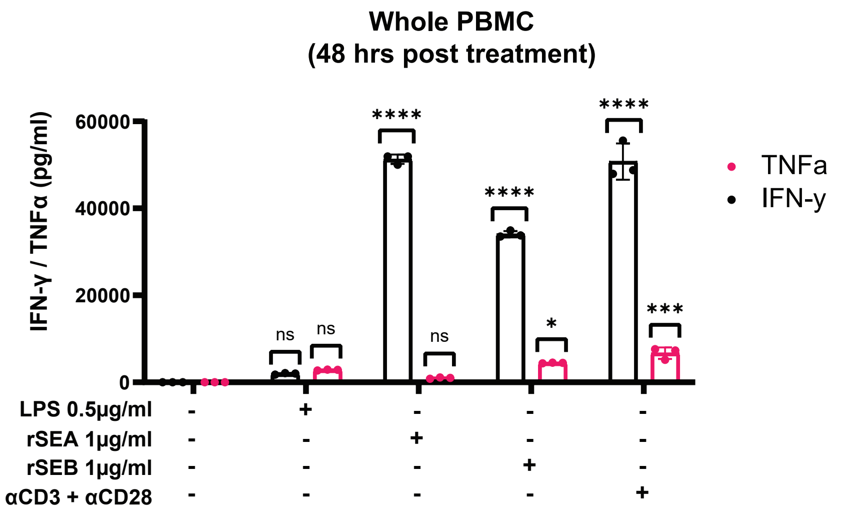


B.

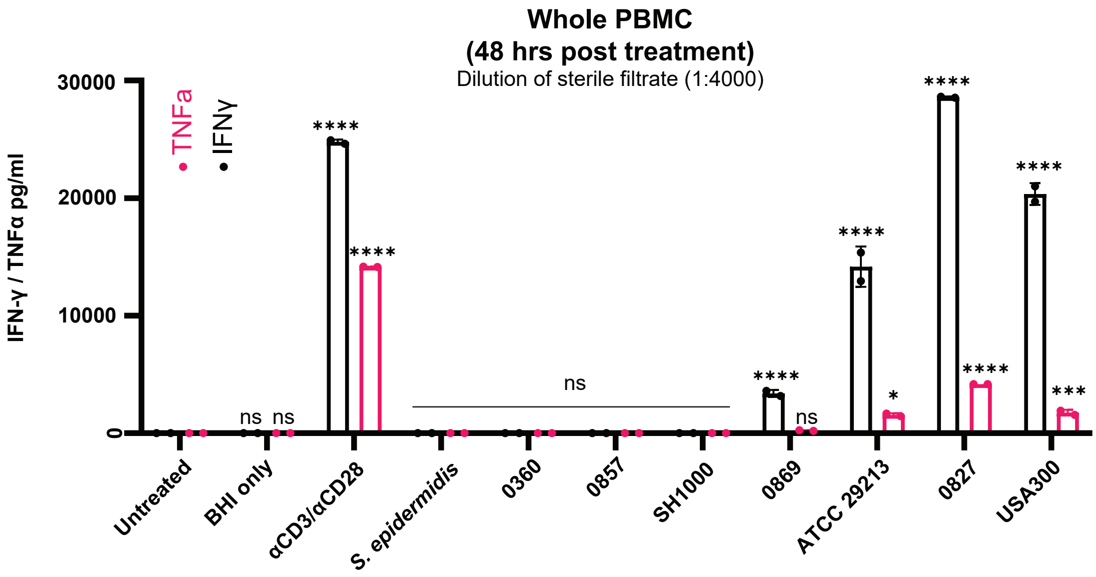


C.

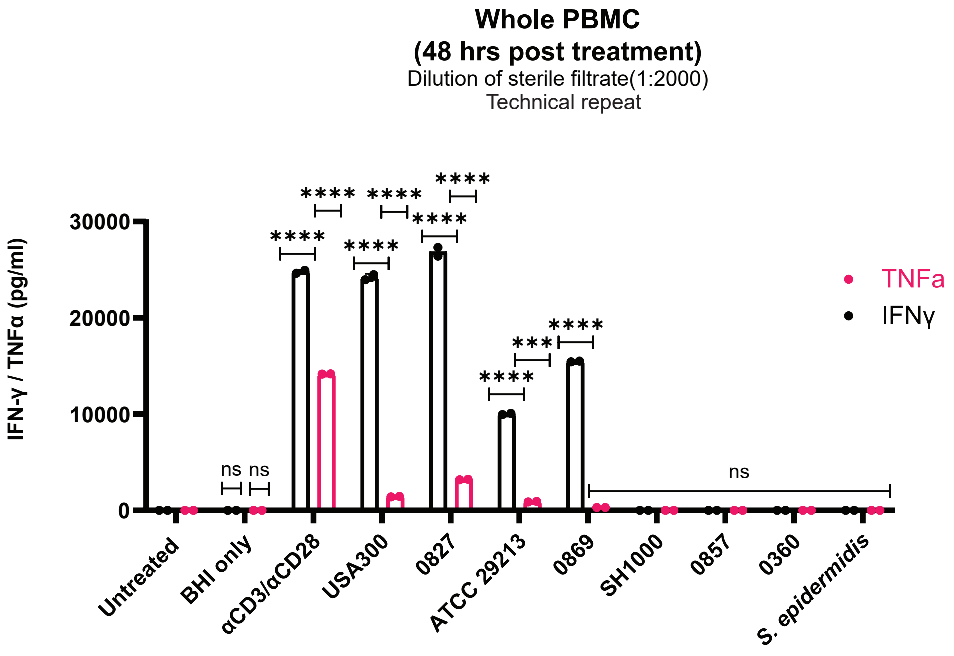

### Fig S10 (For Fig 4)

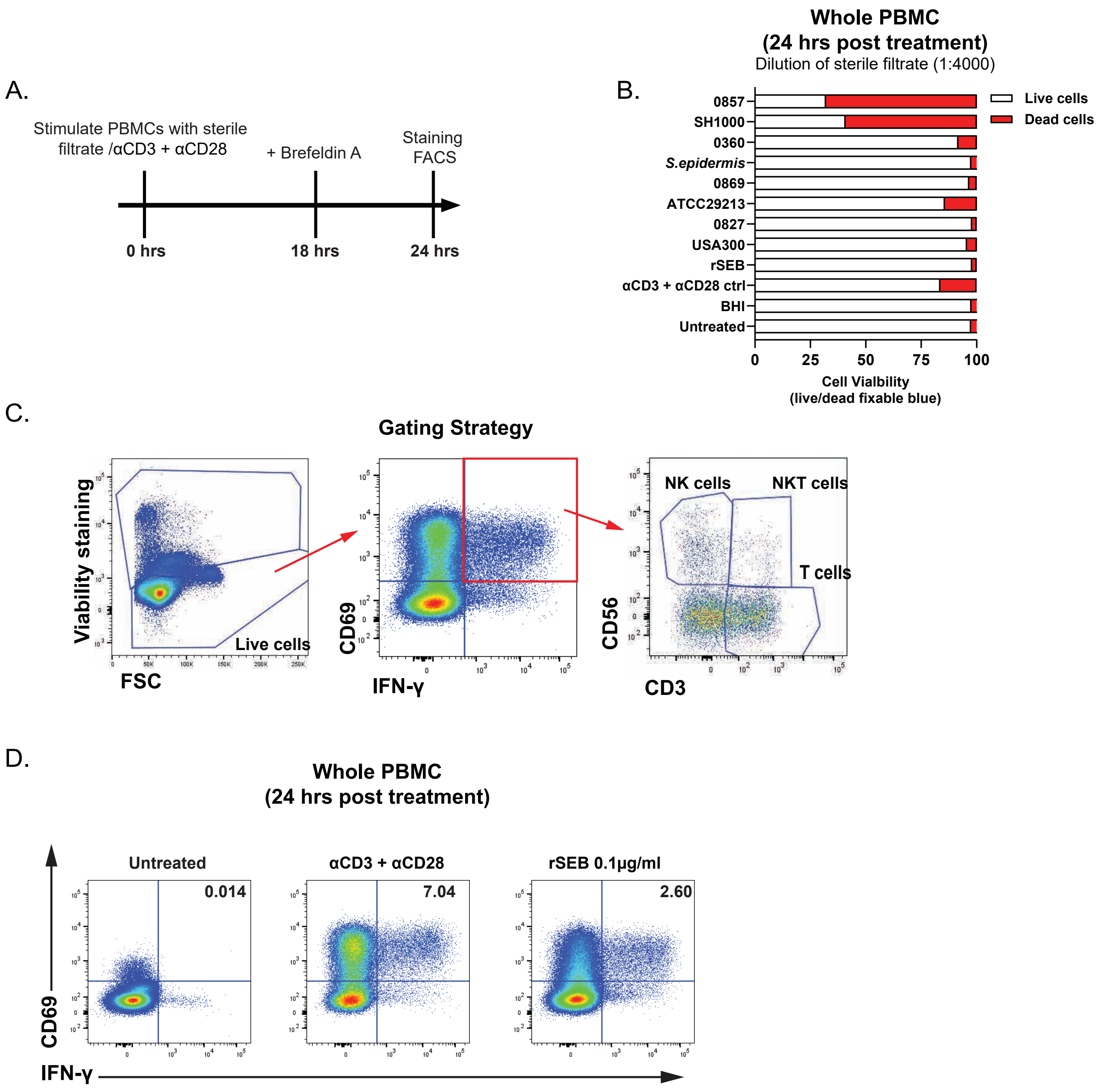

### Fig S11 (For Fig 5)

A.

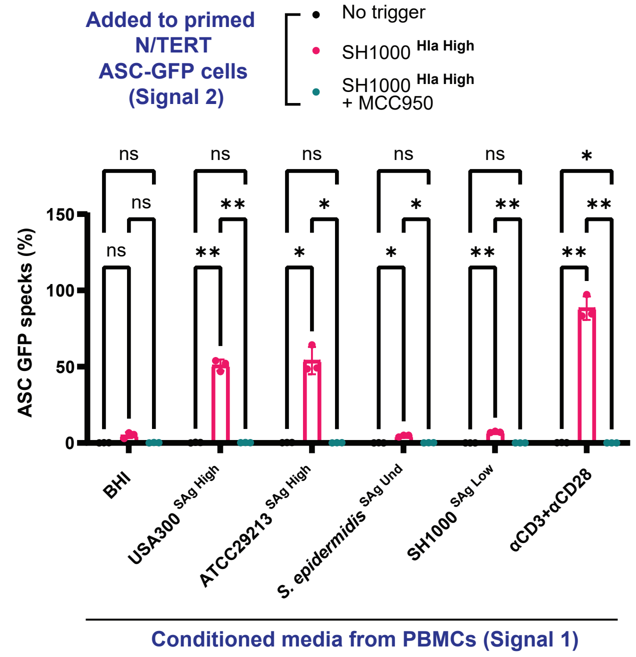


C.

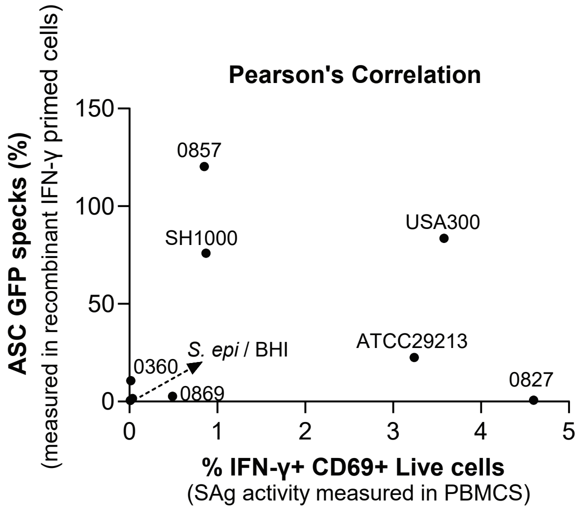


B.

| | Hla | SAg |
|---|---|---|
| SH1000 | High | Low |
| USA300 (MRSA) | High | High |

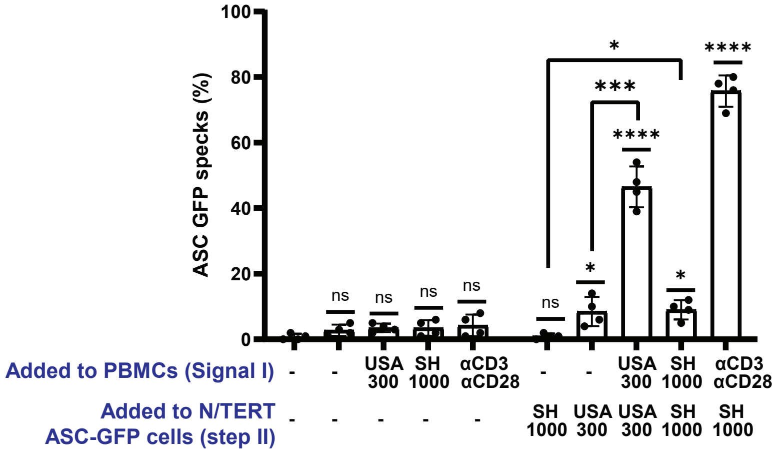

### Fig S12 (For Fig 5)

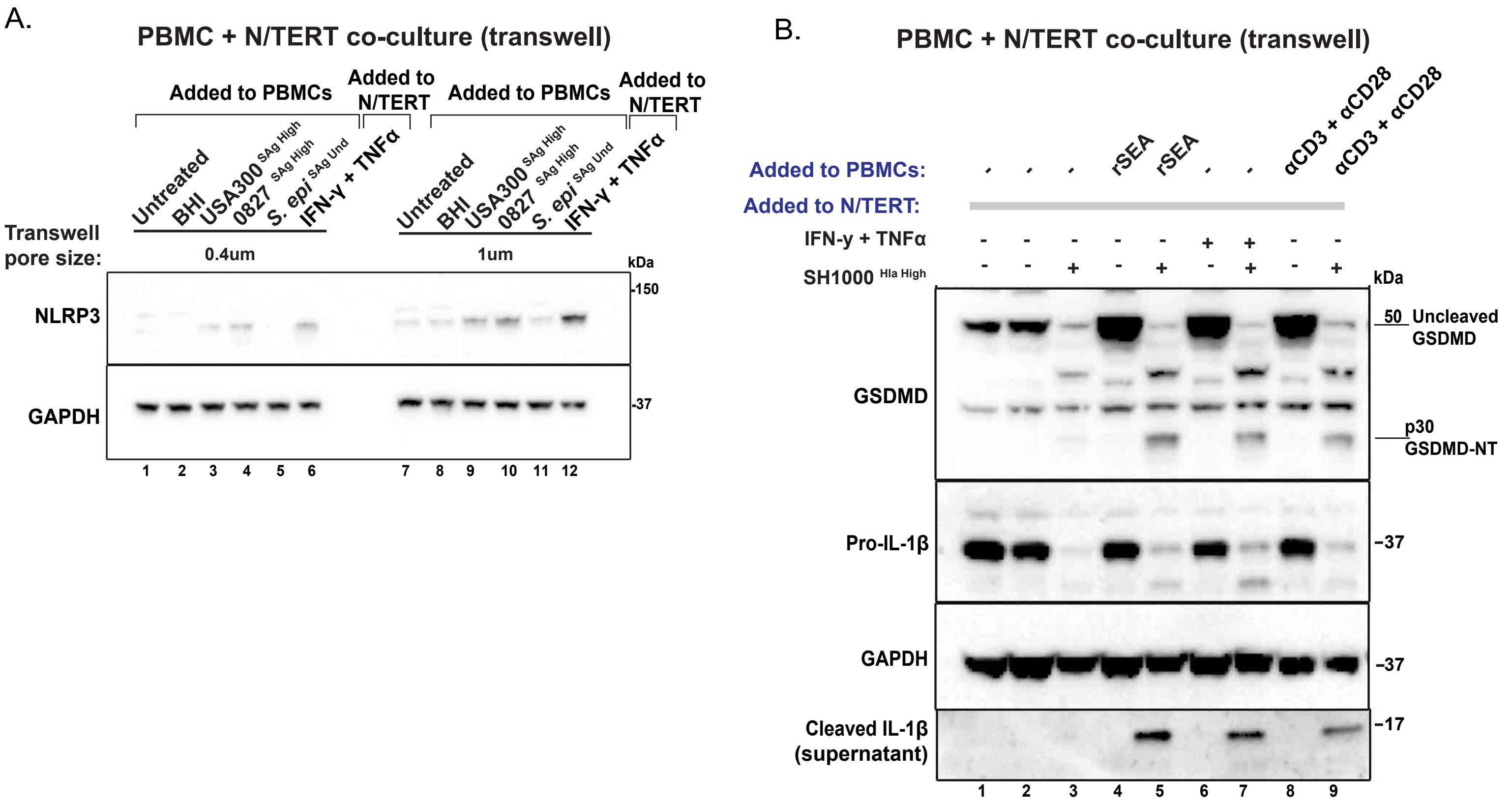
